# A programmable genetic platform for engineering noninvasive biosensors

**DOI:** 10.1101/2025.09.07.674790

**Authors:** Asish N. Chacko, Kaamini M. Dhanabalan, Jinyang Wan, Roy Chien, Nolan T. Anderson, Binzhi Xu, Katie Pham, Ritu Tiwari, Arnab Mukherjee

**Affiliations:** Department of Chemistry, University of California, Santa Barbara, CA 93106, USA; Department of Chemical Engineering, University of California, Santa Barbara, CA 93106, USA; Department of Ecology, Evolution, and Marine Biology, University of California, Santa Barbara, CA 93106, USA; Interdisciplinary Program in Quantitative Biosciences, University of California, Santa Barbara, CA 93106, USA; Department of Molecular, Cellular, and Developmental Biology, University of California, Santa Barbara, CA 93106, USA; Department of Diagnostic and Biomedical Sciences, School of Dentistry, University of Texas Health Science Center at Houston, TX 77054, USA

**Keywords:** aquaporins, biosensors, MRI, reporter genes, protease circuits

## Abstract

Creating genetic sensors for noninvasive visualization of biological activities in deep, optically opaque tissues holds immense potential for basic research and the development of genetic and cell-based therapies. MRI stands out among deep-tissue imaging methods for its ability to generate high-resolution images without ionizing radiation. However, the adoption of MRI as a mainstream biomolecular technology has been hindered by the lack of adaptable methods to link molecular events with genetically encodable MRI contrast. To address this challenge, we introduce universal reporter circuit-based activatable sensors (URCAS), a highly programmable platform for the systematic creation of genetic sensors for MRI. In developing URCAS, we engineered protease-activatable MRI reporters using two distinct approaches: protein stabilization and subcellular trafficking. We established the applicability of URCAS in five diverse mammalian cell types and showcased its versatility by assembling a toolkit of genetic sensors for viral proteins, small-molecule drugs, logic gates, protein-protein interactions, and calcium, without requiring new customization for each target. Our findings suggest that URCAS provides a modular, programmable platform for streamlining the development of noninvasive, nonionizing, and genetically encoded sensors for biomedical research and in vivo diagnostics.

## Introduction

Genetically encoded sensors have transformed our ability to visualize biological phenomena within living cells by connecting molecular-level events to measurable signals, most commonly fluorescence^1,2^. The rapid expansion of the fluorescent genetic indicator toolkit, now encompassing hundreds of sensors capable of monitoring a wide spectrum of cellular activities, has been driven by the development of broadly adaptable biomolecular sensor-engineering strategies^3,4^. For instance, circular protein permutation^5,6^ and the evolution of modular protein pairs^7–9^ for Förster resonance energy transfer (FRET) offer universal and scalable strategies for the systematic engineering of genetic indicators based on conserved reporter scaffolds, such as the green fluorescent protein (GFP). Despite their transformative impact on cellular imaging, the utility of fluorescent indicators in whole-organism imaging is constrained by the inherent limitations of light scattering and absorption in biological tissues^10–12^. This limitation presents a significant barrier to elucidating physiological and pathological mechanisms in vivo, monitoring disease progression, and tracking cell- and gene-based therapies. Overcoming this limitation requires a conceptual shift in bioimaging — one that leverages the strengths (viz., scalability, versatility) of genetic sensor-engineering principles, while integrating the ability to penetrate deep, optically dense tissues.

Magnetic resonance imaging (MRI) is unparalleled among deep-tissue imaging methods for its ability to generate high-resolution tomographic images with intricate anatomical detail across extensive tissue volumes. Crucially, MRI achieves these benefits without the use of ionizing radiation, instead harnessing tissue-penetrant radiofrequencies to excite water protons polarized in a strong magnetic field. However, as a biomolecular technology, MRI faces a significant limitation when compared to optical methods: the lack of widely adaptable sensor-engineering strategies to connect molecular events with genetically encoded contrast^13^. Consequently, the development of genetic indicators for MRI typically requires resource- and time-intensive protein engineering efforts tailored to each new cellular target. The inherently low throughput of MRI further compounds this challenge, hindering the straightforward application of library-based protein engineering approaches such as directed evolution. Consequently, the repertoire of genetic indicators for MRI remains limited, encompassing a mere 4-5 analytes (to our knowledge)^14–21^. This paucity is particularly striking when contrasted with the hundreds of genetic indicators available for fluorescence imaging, especially considering that the first reporters for both modalities emerged around the same time^22–26^.

Recent studies have demonstrated that aquaporins, which are water channels, can genetically encode MRI contrast by safely enabling the rapid exchange of water molecules between the intra- and extracellular compartments^27–34^. This mechanism allows cells engineered to overexpress aquaporins to be detected using standard MRI techniques that monitor the movement of water molecules^35,36^. One such technique, diffusion-weighted imaging, employs pulsed magnetic gradients separated by a user-defined time interval to make MRI signal intensity sensitive to the thermal displacement of water molecules within that interval. Cellular water diffusivity (D) can be quantified using the equation D ∝−log(intensity)/b, where b represents the magnitude of diffusion weighting due to magnetic gradients. Through diffusion-weighted imaging, aquaporin-expressing cells can be easily identified by their higher diffusivity than wild-type cells, which typically show limited endogenous expression of water channels.

Aquaporins, with their compact genetic footprint (single gene, < 1 kb), high sensitivity, and autonomous contrast mechanism, offer an ideal scaffold for expanding the repertoire of MRI-compatible genetic sensors. To explore this concept, we engineered aquaporins to be activated by viral proteases, which are orthogonal to the mammalian proteome and offer well-established programmable units for detecting a diverse array of biomolecular inputs through protein engineering and synthetic protease-controlled circuits^37–42^. We established two distinct mechanisms to achieve protease modulation of aquaporin-dependent MRI signals: stabilization of aquaporins and their intracellular trafficking from the endoplasmic reticulum to the cell membrane, leading to a programmable protease-based bioengineering platform for constructing MRI sensors, termed “universal reporter circuit-based activatable sensors” (URCAS). Through systematic linker engineering, we optimized URCAS to achieve high fold-activation, and demonstrated its modularity, combinatorial capabilities, and functionality across diverse cell types. We successfully applied URCAS to engineer MRI-based genetic indicators for pathogenic enzymes, small-molecule drugs, protein-protein interactions, biological logic gates, and calcium using a protease as a reconfigurable “Lego-like” common core for target detection, thus eliminating the need for case-specific engineering of distinct biosensing mechanisms.

## Results and Discussion

### Engineering DD-URCAS based on proteolytic rescue of destabilized aquaporin

To engineer protease-activatable aquaporins, we fused the human aquaporin-1 isoform (hAqp1) to destabilizing domains (DDs) separated by the canonical substrate for a variant of the NIa tobacco etch virus protease (TEVP) engineered for high catalytic activity^43^. We hypothesized that, in the absence of TEVP activity, the hAqp1-DD fusion protein is rapidly degraded (MRI off-state), whereas protease activity cleaves the DD to restore hAqp1-based diffusivity signals (on-state) (Fig. 1A). To identify the optimal approach for modulating hAqp1 stability, we fused hAqp1 with engineered DDs derived from the estrogen receptor ligand binding domain^44^, the mammalian FKBP12 prolyl isomerase^45^, and a natural degron comprising the first 80 amino acids of the yeast transcription factor, Rpn4^46^. We placed the fusion constructs under the control of the human EF1α promoter to drive constitutive expression in Chinese hamster ovary (CHO) cell lines. We co-expressed TEVP from an inducible minimal CMV promoter, thereby allowing us to measure hAqp1 signals in both on- and off-states within the same cellular chassis. After independently confirming TEVP activity in these cell lines using a GFP-based sensor^47^ (Fig. S1), we used flow cytometry (Fig. S2) to sort doubly transduced cells (based on co-transcribed fluorescent markers) and performed live-cell diffusion-weighted imaging (Fig. 1B). As expected, TEVP expression alone did not alter the baseline water diffusivity of non-engineered cells or cells expressing hAqp1 without a fused DD (Fig. S3). However, all the hAqp1-DD constructs exhibited significant TEVP-triggered changes in diffusivity, with the largest fold-increase (92 ± 4%, mean ± s.e.m) observed for hAqp1 fused to the FKBP12-DD (Fig. 1C, S4). Western blot analysis of the membrane extracts from these cells revealed a prominent band corresponding to the molecular weight of hAqp1 (30 kDa) under TEVP-induced conditions, (Fig. 1D, S5A). In contrast, when TEVP was turned off, we observed a distinctly weaker slower-migrating band, likely indicative of the uncleaved hAqp1-FKBP12-DD fusion (45 kDa) that had not been completely degraded. Corroborating the immunoblotting results, confocal imaging demonstrated distinct cell surface localization of hAqp1 exclusively in the presence of TEVP expression (Fig. 1E).

**Figure 1.**
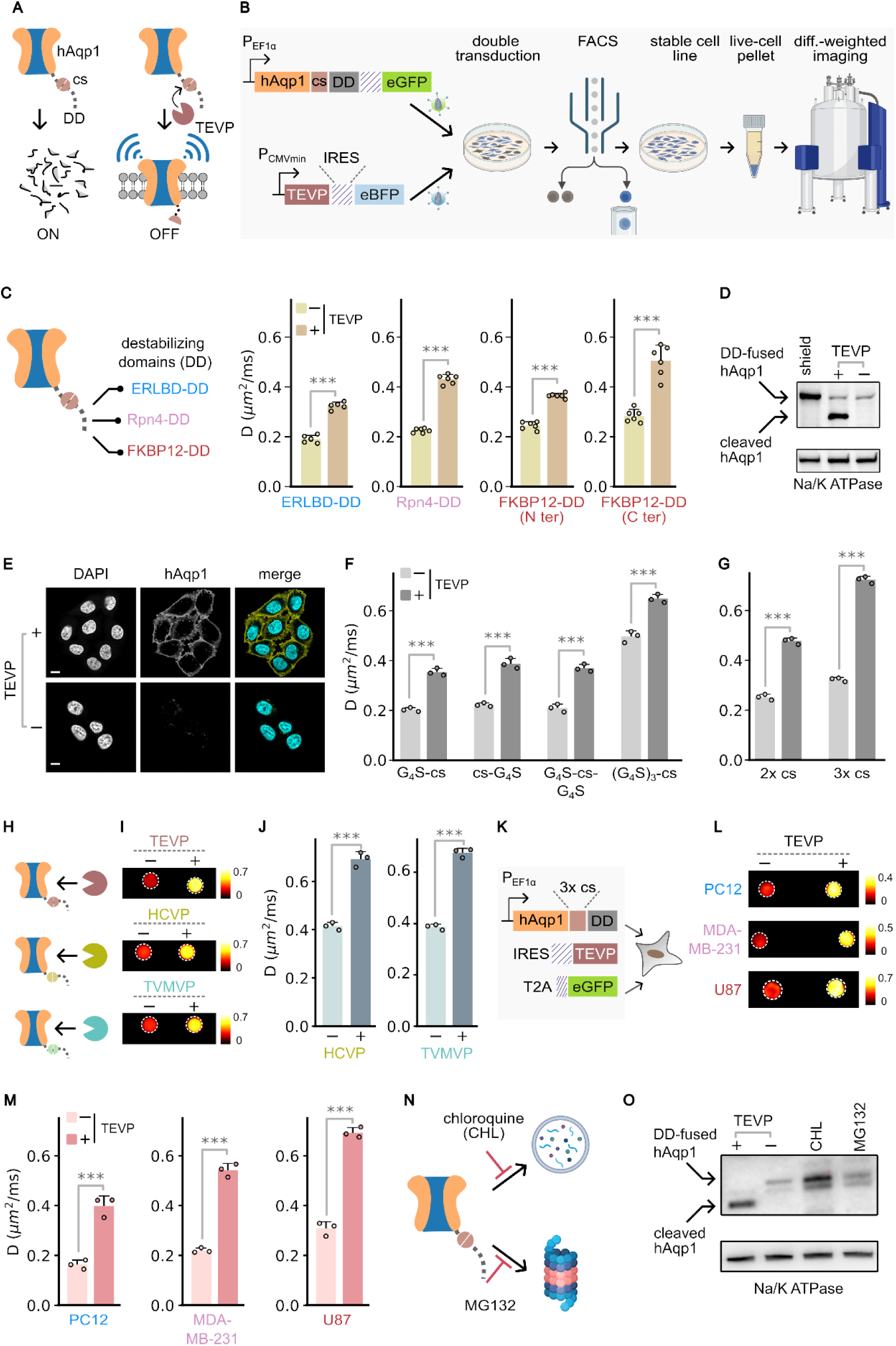
Design, optimization, and evaluation of DD-URCAS. (A) Schematic representation of the mechanism by which hAqp1-based MRI signals, quantified in terms of diffusivity (D, µm^2^/ms), are activated via proteolytic cleavage of a destabilizing domain (DD). (B) Experimental approach for constructing and evaluating protease-activatable aquaporins involves the viral transduction of TEVP and hAqp1 tagged with TEVP-cleavable DD(s) in cell lines, sorting for doubly transduced cells based on co-transcribed fluorescent reporters, and measuring the TEVP-induced change in diffusivities of live-cell pellets using diffusion-weighted MRI. (C) Diffusivities of CHO cells expressing hAqp1 constructs fused with various TEVP-cleavable DD(s), measured both in the absence and presence of TEVP expression. (D) Representative Western blot of membrane extracts from CHO cells engineered to express hAqp1 fused at its C-terminus to the TEVP-cleavable FKBP12-DD, with and without TEVP induction. Membrane lysates from cells treated with shield-1, which stabilizes the FKBP12-DD, are included as a positive control. (E) Representative confocal images of CHO cells expressing hAqp1 fused at its C-terminus to the TEVP-cleavable FKBP12-DD, with and without TEVP induction. Scale bar is 10 µm. (F) Diffusivities of CHO cells expressing hAqp1 tagged at the C-terminus with the FKBP12-DD with Gly-Ser spacers flanking the TEVP cleavage site (cs), measured in the absence and presence of TEVP expression. (G) Diffusivities of CHO cells harboring hAqp1 with multiple TEVP cleavage sites inserted before the FKBP12-DD, obtained with and without TEVP expression. The optimized DD-URCAS configuration harbors three TEVP cut sites and exhibits the largest TEVP-driven fold-change. (H) The modularity of DD-URCAS facilitates straightforward adaptation for multiple proteases. (I) Representative diffusion maps of CHO cell pellets expressing DD-URCAS with and without induction of the corresponding protease. Diffusion maps are depicted using pseudo-color by assigning diffusivity values to a ‘redhot’ color scale. The minimum and maximum diffusivity values (in µm^2^/ms) are specified at the lower and upper edges of the accompanying color bars. (J) Diffusivities of CHO cell pellets expressing DD-URCAS with and without induction of the corresponding protease. (K) Polycistronic genetic constructs for implementing DD-URCAS in different cell lines; eGFP serves as a marker to enrich stably transduced cells by flow sorting. (L) Representative diffusion maps of pellets of various cell types expressing DD-URCAS with and without TEVP expression. Diffusion maps are depicted using pseudo-color as in (I). (M) Diffusivities of pellets of various cell types expressing DD-URCAS with and without TEVP expression. (N) Schematic illustrating the putative degradation pathways of DD-URCAS, and their inhibition by small-molecule modulators, chloroquine and MG132. (O) Representative Western blot of whole-cell lysates prepared from cells expressing DD-URCAS, with or without TEVP expression, and in the presence of the lysosomal inhibitor, chloroquine or the proteasomal inhibitor, MG132. Error bars represent standard deviation from n ≥ 3 measurements. Statistical significance is denoted by *, which indicates P < 0.05; ** denoting P < 0.01; *** denoting P < 0.001; while n.s. indicates P ≥ 0.05.

The FKBP12-DD can be stabilized by a small molecule known as shield-1^45^. When cells expressing hAqp1 fused to the FKBP12-DD were incubated with shield-1, there was a more significant increase in diffusivity (166 ± 15%) compared to what was achieved through TEVP activity (Fig. S5B). Consequently, we pursued two strategies to test whether the TEVP-triggered MRI response could be enhanced further: incorporating flexible spacers around the cleavage site and introducing multiple cleavage sites. While insertion of short Gly-Ser spacers had no notable improvement, a longer 15-residue Gly-Ser linker increased both baseline and TEVP-induced diffusivities, thereby reducing the overall response (Fig. 1F). Conversely, increasing the number of cleavage sites to three resulted in a diffusivity change (124 ± 3%) exceeding that observed with the single site construct (Fig. 1G, S6A). Both single- and triple cleavage site constructs exhibited similar levels of hAqp1 transcription as measured by quantitative RT-PCR, thereby ruling out variations in transduction efficiency as a potential source of the improved response (Fig. S6B). This phenomenon was also observed in hAqp1 fusion with the Rpn4-based degron, wherein the incorporation of triple cleavage sites enhanced the response amplitude compared to a single site, presumably by increasing the statistical rebinding of TEVP to its substrate, analogous to the avidity effect in multivalent binding interactions^48^ (Fig. S6C). Based on these observations, we selected hAqp1 tagged C terminally with the FKBP12-DD and containing upto three protease cut sites as our optimal configuration for the DD-URCAS sensor engineering platform.

The modular architecture of DD-URCAS is expected to facilitate its adaptation to other proteases by simply substituting the TEVP cleavage site with the substrate specific to the target protease. To explore this possibility, we replaced the TEVP substrate with those for the tobacco vein mottling virus protease (TVMVP) and the hepatitis C virus NS3 protease (HCVP) and subsequently expressed the destabilized hAqp1 reporter in cells engineered to produce the respective protease. Upon induction, both TVMVP- and HCVP-based DD-URCAS cells demonstrated notable diffusivity changes of 75 ± 3% and 67 ± 4%, respectively (Fig. 1H-J, S7). To confirm the versatility of the DD-URCAS system across different cell types, we evaluated its efficacy in a dopaminergic neural cell line (PC12), triple-negative breast cancer cell line (MDA-MB-231), and glioblastoma cell line (U87). Each cell line was engineered to express hAqp1-FKBP12-DD, with or without TEVP co-transcribed via IRES (Fig. 1K). We observed substantial changes in MRI signals, ranging from 125 ± 10% to 147 ± 8% across all cell types (Fig. 1L-M, S8).

To further explore the degradation mechanism in DD-URCAS, we exposed CHO cells in the TEVP-off state to the small-molecule drugs chloroquine and MG132, which inhibit the lysosomal^49^ and proteasomal^50^ pathways for protein degradation, respectively (Fig. 1N). Western blot analysis of extracts from chloroquine-treated cells revealed a distinct band corresponding to intact hAqp1-FKBP12-DD fusion protein (Fig. 1O, S5C). In contrast, a fainter-band was observed in cells that had been treated with MG132, suggesting that hAqp1-FKBP12-DD is largely degraded in these cells. Notably, while chloroquine treatment inhibited hAqp1 degradation, it did not enhance its diffusivity (Fig. S9A), suggesting that although lysosomal degradation was blocked, hAqp1-FKBP12-DD was not expressed durably in the plasma membrane to elicit a significant change in diffusivity. We confirmed this through confocal imaging of chloroquine-treated cells, which showed a clear accumulation of hAqp1 in vesicular endosomal structures but not in the cell membrane (Fig. S9B). Collectively, these findings provide mechanistic evidence that FKBP12-DD tagged hAqp1 is primarily removed via the lysosomal pathway and that excising the DD allows for stable surface expression of hAqp1, thereby restoring diffusivity.

### Programming DD-URCAS for biosensing applications

Small-molecule drugs that modulate protease activity are coveted for both clinical applications, such as antiviral therapy^51,52^, and for regulating protease-based synthetic gene circuits^37,38,53–55,56^. Therefore, we first explored the potential of DD-URCAS to detect various protease-modulating small-molecule drugs. Our initial focus was on trimethoprim, an antibiotic that has also been repurposed to engineer chemically switchable biologics for numerous gene and cell-based therapies^57–63^. To make TEVP responsive to trimethoprim, we fused TEVP with a DD derived from E. coli dihydrofolate reductase (ecDHFR)^64,65^. The resulting TEVP-ecDHFR fusion is unstable, but can be stabilized by adding trimethoprim, which binds to and stabilizes ecDHFR (Fig. 2A). As anticipated, DD-URCAS cells engineered to co-express ecDHFR-tagged TEVP exhibited diffusivity values near baseline (Fig. 2B); however, treatment with trimethoprim led to a dose-dependent increase in diffusivity (Fig. 2C), with diffusivity signals fully restored at 10 µM trimethoprim (Fig. 2B). Subsequently, we proceeded to image small-molecule inhibitors (Fig. 2D), selecting the clinically relevant hepatitis C virus NS3 protease (HCVP) as our target. Incubating the HCVP-based DD-URCAS cells with clinically approved inhibitors, boceprevir^66^ and telaprevir^67^, reduced the MRI signals to levels observed in cells without HCVP expression (Fig. 2E,F).

**Figure 2.**
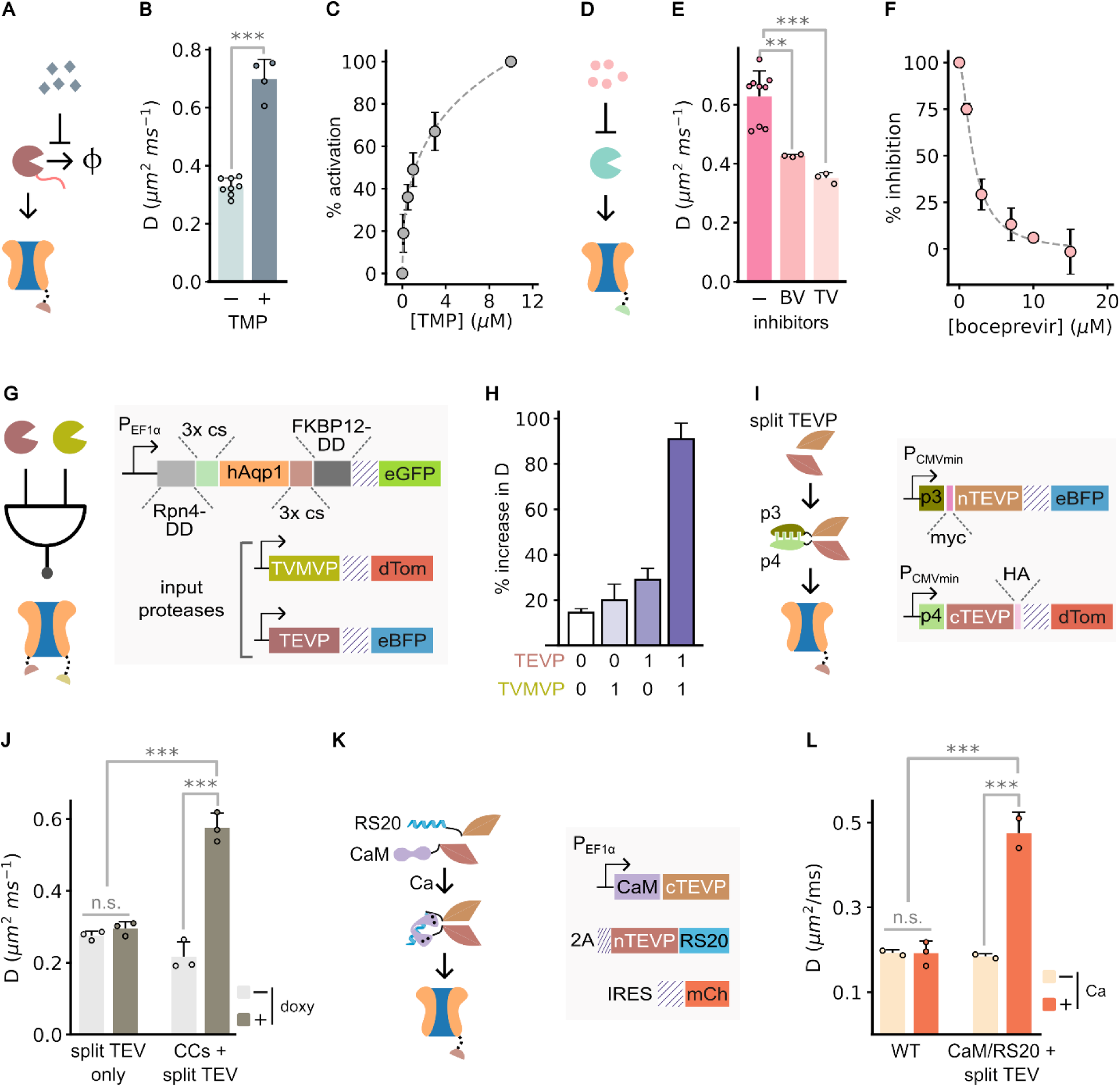
Programmable biosensor design based on DD-URCAS. (A) Schematic representation illustrating the integration of DD-URCAS with destabilized proteases for the detection of the small-molecule drug trimethoprim (TMP). (B) Quantification of diffusivities in CHO cells engineered to express the TMP-sensing DD-URCAS circuit as depicted in (A), acquired in the presence and absence of TMP treatment for 24 h. (C) Dose-response analysis of the TMP-sensing DD-URCAS expressed in CHO cells incubated with varying concentrations of TMP for 24 h. (D) Schematic depiction of the application of DD-URCAS for the detection of small-molecule inhibitors of protease activity. (E) Diffusivities of CHO cells engineered to express HCVP-modulated DD-URCAS, with and without treatment using antiviral HCVP inhibitors: boceprevir (BV) or telaprevir (TV) for 24 h. (F) Dose-response analysis of the HCVP-based DD-URCAS in CHO cells exposed to varying concentrations of boceprevir for 24 h. (G) A schematic illustrating the detection of a protease-based 2-input biological AND gate using DD-URCAS. The corresponding genetic constructs for introducing the 2-input AND gates and DD-URCAS are shown alongside. (H) Diffusivities of CHO cells engineered to express the protease-based AND gate and corresponding DD-URCAS detector, in the presence and absence of the respective protease inputs. (I) Integration of DD-URCAS with split protease technology to image protein-protein interactions. The corresponding genetic constructs are shown alongside. (J) Diffusivities of CHO cells expressing the split TEVP-based DD-URCAS for detecting protein-protein interaction. Control measurements are obtained in cells expressing split TEVP fragments without the fused coiled-coil peptides. (K) Schematic illustrating the engineering of split TEVP-based DD-URCAS for calcium imaging and the corresponding genetic constructs. The corresponding genetic constructs for implementing the calcium recorder are shown alongside. (L) Diffusivities of CHO cells expressing the calcium-sensing DD-URCAS circuit, measured with and without incubation with a calcium ionophore, ionomycin. Control measurements are obtained in identically treated wild-type cells lacking the sensor.. Error bars represent standard deviation from n ≥ 3 measurements. Statistical significance is denoted by *, which indicates P < 0.05; ** denoting P < 0.01; *** denoting P < 0.001; while n.s. indicates P ≥ 0.05.

Given that the DD-URCAS paradigm generalizes in a straightforward manner to orthogonal proteases, we investigated its potential for detecting biological logic gates, which are valuable motifs for designing combinatorial gene circuits controlled by multiple inputs (e.g., cell state classifiers). As a proof-of-concept, we aimed to develop a sensor for detecting AND logic, in which the simultaneous activity of two protease inputs is necessary to generate an MRI signal. To adapt DD-URCAS for sensing protease-based AND gates, we needed hAqp1 to be modulated concurrently by two orthogonal proteases. Therefore, in addition to the FKBP12-DD at the C-terminus of hAqp1, we fused a second degron derived from Rpn4 at the N-terminus (Fig. 2G). We inserted cut sites for TVMVP and TEVP upstream of each DD, and introduced the resulting sensor into CHO cells co-expressing TEVP, TVMVP, or both proteases (Fig. 2G). As a negative control, we generated CHO cells expressing HCVP, whose recognition site is distinct to both TEVP and TVMVP. As expected, the simultaneous activity of both TEVP and TVMVP was necessary to trigger a significant change in MRI signals, which exceeded the response observed with no or only one protease by at least 3-fold (P-value < 10^-3^) (Fig. 2H).

To demonstrate the versatility of DD-URCAS for biosensor engineering, we integrated this method with split protease technology to create additional MRI-based indicators for detecting biomolecular binding and intracellular calcium. In this strategy, TEVP is divided into amino acids T119, producing two fragments: N-TEVP consisting of amino acids 1-119 and C-TEVP comprising amino acids 120-243. This design deactivates TEVP by separating the catalytic triad, forming the foundation of a broadly applicable sensing mechanism in which TEVP is conditionally reactivated by inputs that promote the assembly of split fragments^68^ (Fig. 2I). In the first application, we engineered sensors to detect binding between a pair of coiled-coils using the well-known heterodimerizers p3 and p4^69^. We attached p3 and p4 at the N-terminus of each TEVP fragment (Fig. 2I) and expressed them in CHO cells. Control cells were engineered to express split TEVP fragments without the fused dimerizers. Expressing individual TEVP fragments without the p3/p4 pair did not trigger any diffusivity change. However, when the TEVP fragments were fused to heterodimerizing peptides, protease activity was restored, resulting in 165 ± 7% change in diffusion signals (Fig. 2J, S10A). To adapt the split TEVP-based DD-URCAS for calcium imaging, we fused N-TEVP to calmodulin, and C-TEVP to a myosin light chain kinase peptide, RS20 (Fig. 2K). Calmodulin binds to RS20 in the presence of calcium, forming the basis for most fluorescent calcium biosensors^70^. We expressed the calmodulin/RS20-fused split TEVP-URCAS (Fig. 2K) in CHO cells and induced cytosolic calcium entry by treating the cells with the calcium ionophore ionomycin. A significant 157 ± 7% increase in MRI signals was observed in response to ionomycin treatment in the DD-URCAS cells but not in similarly treated wild-type cells (Fig. 2L, S10B).

### Engineering ER-URCAS based on proteolytic removal of an ER retention tag

After demonstrating the ability to modulate MRI signals by proteolytically controlling hAqp1 degradation, we investigated whether targeted proteolysis can be used to develop hAqp1-based indicators without having to destabilize the hAqp1 reporter. Since hAqp1 must enter the ER before localizing to the plasma membrane, we considered whether cells could be engineered to load a pool of pre-formed hAqp1 in the ER (where it cannot generate MRI contrast) and release this pool only in response to specific sensory inputs via targeted proteolytic cleavage. Proteins residing in the ER or cycling between the ER and post-ER compartments, such as the Golgi, are often marked with retention signal peptides (e.g., KKXX peptides) at their C-terminus^71^. Therefore, tagging hAqp1 C-terminally with a retention signal separated by a protease cut site could provide a mechanism for the controlled release of pre-formed hAqp1 from the ER lumen through proteolytic excision of the retention tag (Fig. 3A). This approach has been recently demonstrated for the controlled secretion of protein-based therapeutics^72–74^. Ideally, a trafficking-based modulation approach could offer specific advantages over the degradation-based approach by insulating sensor operation from innate degradation pathways and reducing the genetic footprint (12 vs. 321 base pairs for the trafficking- and DD-based designs).

**Figure 3.**
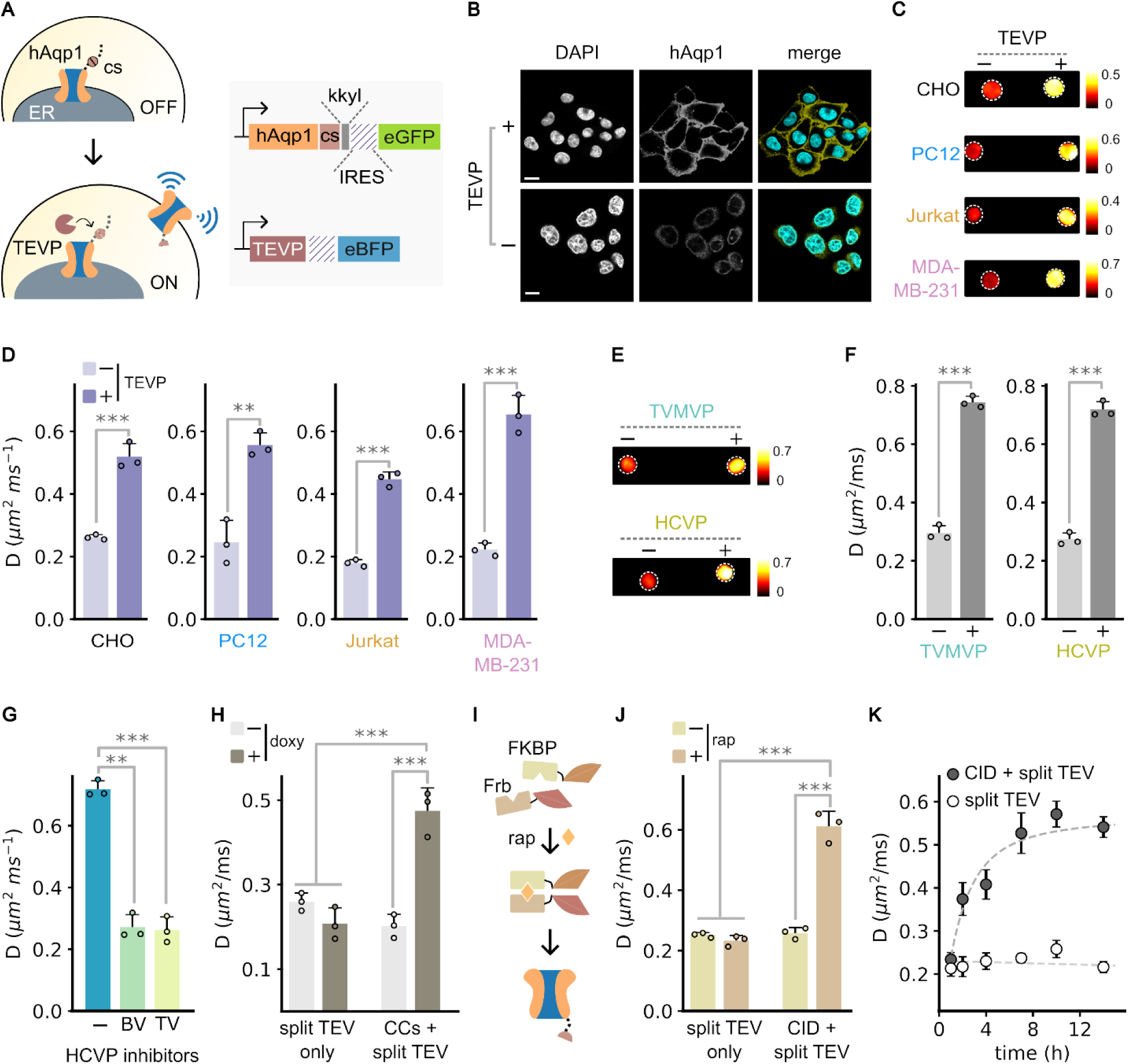
Programmable biosensor design based on ER-URCAS. (A) Schematic representation of the mechanism by which hAqp1-based MRI signals quantified in terms of diffusivity (D, µm^2^/ms), are activated via proteolytic cleavage of an ER retention tag (KKYL). Genetic constructs encoding ER-URCAS are depicted alongside. (B) Representative confocal images of CHO cells transduced with ER-URCAS, acquired with and without TEVP expression. (C) Representative diffusion maps of various cell types expressing ER-URCAS, with and without TEVP expression. Diffusion maps are depicted using pseudo-color by assigning diffusivity values to a ‘redhot’ color scale. The minimum and maximum diffusivity values (in µm^2^/ms) are specified at the lower and upper edges of the accompanying color bars. (D) Diffusivities of pellets of various cell types expressing ER-URCAS, with and without TEVP expression. (E) Representative diffusion maps CHO cell pellets expressing ER-URCAS with and without induction of the corresponding protease. . Diffusion maps are depicted using pseudo-color as in (C). (F) Diffusivities of CHO cell pellets expressing ER-URCAS with and without induction of the corresponding protease. (G) Diffusivities of CHO cells engineered to express HCVP-modulated ER-URCAS, with and without treatment using antiviral HCVP inhibitors: boceprevir (BV) or telaprevir (TV) for 24 h. (H) Diffusivities of CHO cells expressing the split TEVP-based ER-URCAS sensor for detecting protein-protein interaction. Control measurements are obtained in cells expressing split TEVP fragments without the fused p3/p4 coiled-coils. (I) Schematic illustrating the adaptation of split TEVP-based ER-URCAS sensor to detect chemically induced dimerization of FKBP and Frb induced by rapamycin. (J) Diffusivities of CHO cells expressing split TEVP-based sensor depicted in (J). Control measurements are obtained in cells expressing split TEVP without the fused FKBP and Frb domains. (K) Time-dependent activation of ER-URCAS in CHO cells engineered to express FKBP and Frb-fused split TEVP fragments upon rapamycin induction. Control measurements are obtained in cells expressing split TEVP without the fused FKBP or Frb. Error bars represent standard deviation from n ≥ 3 measurements. Statistical significance is denoted by *, which indicates P < 0.05; ** denoting P < 0.01; *** denoting P < 0.001; while n.s. indicates P ≥ 0.05.

To test this concept, we engineered hAqp1 with a KKYL retention sequence at its C-terminus and inserted three cleavage sites for TEVP, which allowed efficient proteolytic excision in our DD-tagged constructs. We transduced cells to co-express KKYL-tagged hAqp1 in conjunction with doxycycline-inducible TEVP (Fig. 3A) and performed both optical imaging and MRI. Confocal microscopy showed that hAqp1-KKYL was largely localized intracellularly in the absence of TEVP, whereas the induction of TEVP expression relocated hAqp1 to the cell surface (Fig. 3B). Consistent with the microscopy findings, live-cell MRI revealed substantial TEVP-dependent diffusivity signals ranging ranging from 98-195% increase in diffusivity across various cell types, including CHO, PC12, Jurkat, and MDA-MB-231 cells (Fig. 3C,D, S11). Similar to DD-URCAS, the ER-URACS approach was modular, enabling the detection of other proteases simply by swapping the TEVP cut site with the substrate for the desired protease (Fig. 3E,F, S12).

### Programming ER-URCAS for biosensing applications

As with the DD-URCAS, we started by demonstrating the ability of ER-URCAS to detect the small-molecule HCVP inhibitors, boceprevir and telaprevir (Fig. 3G). Subsequently, we extended this approach for imaging protein-protein interactions via split protease technology. First, we showed that ER-URCAS can be used to sense heterodimerization of p3/p4 coiled-coils, generating a larger binding-dependent change in diffusion than observed using DD-URCAS (Fig. 3H, S13). Finally, we applied split TEVP-based ER-URCAS to detect chemically-induced protein binding by fusing N-TEVP and (truncated) C-TEVP respectively to the C-termini of FKBP and Frb (rapamycin-binding domain of mTOR). Binding of FKBP to Frb depends on the small molecule dimerizer, rapamycin (Fig. 3I), allowing us to track ligand-induced protein interactions and also characterize the time-scale of activation of our ER-based sensor. Cells expressing FKBP12 and Frb fusions with the truncated split TEVP system showed minimal diffusion change in the absence of rapamycin, but generated a 136% increase in diffusivity following incubation with rapamycin (Fig. 3J, S13). We measured the responsiveness of the sensor by tracking the increase in diffusion as a function of time following rapamycin treatment. As has been recently shown for synthetic ER-release based protein secretion systems, our ER-based sensor exhibited a rapid increase in diffusivity reaching nearly half-maximum signal change within approximately 2 h (Fig. 3K). We also evaluated an alternative configuration of the FKBP-Frb sensor, employing a split TEVP with truncation of C-terminal residues (C-TEVPΔ219-243), which is known to enhance catalytic activity^75^. Expression of the truncated split TEVP-based ER-URCAS yielded significant background diffusivity signals, even in the absence of rapamycin (Fig. S13), indicating that non-specific protein-protein interactions were likely sufficient to activate the truncated split TEVP construct. Taken together, these results validate that proteolytic excision of a tetrapeptide ER-retention tag can be used to rationally design compact biosensors that are insulated from the host cell’s innate degradation pathways.

## Conclusions

This study introduces URCAS as a versatile bioengineering platform for the systematic and scalable creation of genetic sensors for noninvasive imaging. By harnessing viral proteases to recognize specific inputs and transduce these signals into genetically encoded MRI outputs, URCAS addresses a long-standing challenge in the field - namely, the need to custom-design MRI reporters for each new cellular process and analyte. Instead, URCAS exploits the adaptability of viral proteases, which can be rationally reprogrammed using established molecular engineering techniques to detect user-specified cellular events and biomarkers. This work showcases the versatility and modularity of URCAS through the successful engineering of sensors responsive to distinct proteases, small-molecule modulators, and reconstituted protease activity via destabilized or split variants. The reconfigurable “plug-and-play” architecture positions URCAS as a powerful tool for expanding the scope of biomolecular imaging. For example, URCAS can be adapted for detecting protein kinase activity by fusing split TEVP fragments to kinase-inducible interaction domains^76^, or repurposed for high-throughput screening of compounds that modulate disease-relevant protein-protein interactions^77^, thereby advancing in vivo drug discovery campaigns. Further expansion of URCAS’ capabilities can be achieved by integrating additional protease modulation mechanisms. One such strategy involves the proteolytic release of membrane-tethered proteases^79–82^, which can be used to re-engineer URCAS for imaging cell-surface receptor activation and intercellular communication^83^. The incorporation of multiple proteases and protease-based logic also opens the door to integrating URCAS with artificial protease-based sensing networks^38,55,84^ capable of executing more complex tasks, such as classifying cell fates and detecting disease states with high precision and specificity. These pursuits will directly benefit from the availability of a large library of viral proteases with orthogonal cut sites and varying catalytic rates, which have proven pivotal to the recent expansion of protease-based synthetic biological devices for applications in human health^85,86^.

As a fully genetic system, URCAS holds significant promise for clinical translation, especially in gene- and cell-based therapies. URCAS could be genetically integrated with approved cell- and gene-based therapeutics to longitudinally monitor disease progression and therapeutic outcomes. Furthermore, URCAS’s post-transcriptional activation mode allows for versatile in vivo delivery strategies: DNA-based delivery (as employed in this study) facilitates tissue-specific genetic targeting through viral vectors and cell type-specific promoters, while mRNA-based delivery circumvents the need for nuclear entry, potentially minimizing insertional mutagenesis risks and enabling rapid, tunable expression with dynamic dosing capabilities^87^. Notably, both DD- and ER-based URCAS mechanisms operate independently of transcriptional activation, thus avoiding common limitations such as delayed response times and transcriptional silencing. Together, these features position URCAS as a transformative platform for genetic medicine, whose continued development and expansion is expected to align synergistically with ongoing advancements in therapeutic gene delivery.

As with any emerging biotechnology, URCAS has certain limitations. Firstly, although all URCAS configurations show significant fold-changes (ranging from 67% to 195%), further investigation is needed to understand the factors contributing to variations in response amplitude across different sensor architectures. For instance, some constructs exhibited a higher baseline, likely due to incomplete degradation of destabilized hAqp1. Future research could explore the impact of junctional sequences and protease cut sites at the interface of hAqp1 and the DD (or the ER-localizing tag), which may influence the efficacy of degradation (or ER-sequestration) and the steric accessibility of the protease to its specific cut site. Secondly, while using hAqp1 as a deep-tissue reporter offers several distinct advantages (such as a small genetic footprint, human origin, and autonomous contrast), its diffusion-based mechanism could be a limitation if changes in intrinsic cell membrane permeability (e.g., in apoptotic and necrotic cells) do not specifically induce significant changes in diffusivity. Although MRI techniques have been developed to differentiate hAqp1-based diffusivity changes from those caused by alterations in cellular morphology^33^, the application of these methods in the context of URCAS has yet to be directly evaluated. Alternatively, the URCAS concept can be expanded to develop sensors detectable by other MRI modalities that do not rely on diffusion (e.g., T_1_, T_2_, and CEST reporters)^88–92^, as well as emerging techniques such as biomolecular ultrasound^93–95^. These multimodal sensors can complement hAqp1-based URCAS and provide nonoverlapping capabilities to concurrently monitor multiple parameters. In preliminary work (in preparation), we demonstrated the generalizability of the URCAS concept by extending it to a paramagnetic MRI reporter that does not depend on diffusion to generate contrast. Thirdly, further research is crucial to demonstrate and evaluate the URCAS concept in a wide range of in vivo biomedical applications. In this regard, the in vivo application of URCAS would benefit from numerous published examples using hAqp1 as an MRI reporter in mouse models, including imaging tumor gene expression^27,29^, tracking cancer cell-activated promoters^30^, mapping viral gene delivery through the BBB^31^, and tracing neuronal connectivity in the brain^32^. Fourthly, while the reporter components of URCAS are human-derived, protease controllers are of non-human origin, which presents potential immunogenicity risks. Further testing in animal models is necessary to assess these risks.

The conceptual development of URCAS can be likened to the discovery of versatile mechanisms for engineering genetic sensors that utilize fluorescent reporters. These mechanisms have been instrumental in broadening the scope of optical imaging to encompass a wide range of biological processes and analytes. In a similar vein, URCAS represents a highly programmable biosensor technology poised to enhance both the scope and depth of biomedical imaging in areas beyond the reach of optical indicators.

## Author Contributions

AM and ANC conceived this study and were responsible for designing the research methodology and all experiments. KMD executed the experiments related to the design, construction, and testing of the calcium sensor. JW conducted the experiments concerning the design, construction, and testing of the trimethoprim sensor. ANC, with assistance from RC and KP, performed all other experiments. NTA engineered the TEVP-expressing cell lines. BX provided support with cell culture experiments. Data analysis was carried out by AM and ANC, with contributions from KMD, JW, and RT. The manuscript was prepared by AM and ANC, with input from RT. AM supervised the overall project.

## Data availability

The data that support the findings of this study are readily available from the corresponding author upon reasonable request.

## Materials transfer agreement

All plasmid constructs used in this work can be provided by the corresponding author pending scientific review and a completed material transfer agreement. Requests for plasmids should be submitted to: Arnab Mukherjee.

## Acknowledgements.

This research was supported by the National Institutes of Health (R35-GM133530 and R01-NS128278 to A.M.), the U.S. Army Research Office via the Institute for Collaborative Biotechnologies cooperative agreement W911NF-19-D-0001-0009 (A.M.), and a grant 2024-338493 from the Chan Zuckerberg Initiative DAF, an advised fund of Silicon Valley Community Foundation (A.M.). The research made use of the shared facilities of the Materials Research Science and Engineering Center (MRSEC) at UC Santa Barbara: NSF DMR–2308708. The UC Santa Barbara MRSEC is a member of the Materials Research Facilities Network (www.mrfn.org). We acknowledge the use of the NRI-MCDB Microscopy Facility and the Resonant Scanning Confocal supported by NSF MRI grant 1625770. We thank Dr. Ben Lopez for his outstanding technical assistance with the confocal imaging experiments. We thank Dr. Jerry Hu for assistance with the MRI experiments.

## References

(1) Greenwald, E. C.; Mehta, S.; Zhang, J. Genetically Encoded Fluorescent Biosensors Illuminate the Spatiotemporal Regulation of Signaling Networks. Chem. Rev. 2018, 118 (24), 11707–11794. 10.1021/acs.chemrev.8b00333.

(2) Yeh, H.-W.; Ai, H.-W. Development and Applications of Bioluminescent and Chemiluminescent Reporters and Biosensors. Annu. Rev. Anal. Chem. 2019, 12 (1), 129–150. 10.1146/annurev-anchem-061318-115027.

(3) Sanford, L.; Palmer, A. Recent Advances in Development of Genetically Encoded Fluorescent Sensors. In Methods in Enzymology; Elsevier, 2017; Vol. 589, pp 1–49. 10.1016/bs.mie.2017.01.019.

(4) Germond, A.; Fujita, H.; Ichimura, T.; Watanabe, T. M. Design and Development of Genetically Encoded Fluorescent Sensors to Monitor Intracellular Chemical and Physical Parameters. Biophys. Rev. 2016, 8 (2), 121–138. 10.1007/s12551-016-0195-9.

(5) Topell, S.; Hennecke, J.; Glockshuber, R. Circularly Permuted Variants of the Green Fluorescent Protein. FEBS Lett. 1999, 457 (2), 283–289. 10.1016/S0014-5793(99)01044-3.

(6) Anderson, N. T.; Xie, J. S.; Chacko, A. N.; Liu, V. L.; Fan, K.-C.; Mukherjee, A. Rational Design of a Circularly Permuted Flavin-Based Fluorescent Protein. *Chembiochem Eur*. J. Chem. Biol. 2024, 25 (9), e202300814. 10.1002/cbic.202300814.

(7) Ai, H.; Henderson, J. N.; Remington, S. J.; Campbell, R. E. Directed Evolution of a Monomeric, Bright and Photostable Version of *Clavularia* Cyan Fluorescent Protein: Structural Characterization and Applications in Fluorescence Imaging. Biochem. J. 2006, 400 (3), 531–540. 10.1042/BJ20060874.

(8) Miyawaki, A.; Llopis, J.; Heim, R.; McCaffery, J. M.; Adams, J. A.; Ikura, M.; Tsien, R. Y. Fluorescent Indicators for Ca2+based on Green Fluorescent Proteins and Calmodulin. Nature 1997, 388 (6645), 882–887. 10.1038/42264.

(9) Goedhart, J.; Von Stetten, D.; Noirclerc-Savoye, M.; Lelimousin, M.; Joosen, L.; Hink, M. A.; Van Weeren, L.; Gadella, T. W. J.; Royant, A. Structure-Guided Evolution of Cyan Fluorescent Proteins towards a Quantum Yield of 93%. Nat. Commun. 2012, 3 (1), 751. 10.1038/ncomms1738.

(10) Leblond, F.; Davis, S. C.; Valdés, P. A.; Pogue, B. W. Pre-Clinical Whole-Body Fluorescence Imaging: Review of Instruments, Methods and Applications. J. Photochem. Photobiol. B 2010, 98 (1), 77–94. 10.1016/j.jphotobiol.2009.11.007.

(11) Ntziachristos, V. Going Deeper than Microscopy: The Optical Imaging Frontier in Biology. Nat. Methods 2010, 7 (8), 603–614. 10.1038/nmeth.1483.

(12) Helmchen, F.; Denk, W. Deep Tissue Two-Photon Microscopy. Nat. Methods 2005, 2 (12), 932–940. 10.1038/nmeth818.

(13) Yun, J.; Baldini, M.; Chowdhury, R.; Mukherjee, A. Designing Protein-Based Probes for Sensing Biological Analytes with Magnetic Resonance Imaging. Anal. Sens. 2022, 2 (5), e202200019. 10.1002/anse.202200019.

(14) Shapiro, M. G.; Westmeyer, G. G.; Romero, P. A.; Szablowski, J. O.; Küster, B.; Shah, A.; Otey, C. R.; Langer, R.; Arnold, F. H.; Jasanoff, A. Directed Evolution of a Magnetic Resonance Imaging Contrast Agent for Noninvasive Imaging of Dopamine. Nat. Biotechnol. 2010, 28 (3), 264–270. 10.1038/nbt.1609.

(15) Shapiro, M. G.; Szablowski, J. O.; Langer, R.; Jasanoff, A. Protein Nanoparticles Engineered to Sense Kinase Activity in MRI. J. Am. Chem. Soc. 2009, 131 (7), 2484–2486. 10.1021/ja8086938.

(16) Airan, R. D.; Bar-Shir, A.; Liu, G.; Pelled, G.; McMahon, M. T.; Van Zijl, P. C. M.; Bulte, J. W. M.; Gilad, A. A. MRI Biosensor for Protein Kinase A Encoded by a Single Synthetic Gene. Magn. Reson. Med. 2012, 68 (6), 1919–1923. 10.1002/mrm.24483.

(17) Hai, A.; Cai, L. X.; Lee, T.; Lelyveld, V. S.; Jasanoff, A. Molecular fMRI of Serotonin Transport. Neuron 2016, 92 (4), 754–765. 10.1016/j.neuron.2016.09.048.

(18) Ghosh, S.; Li, N.; Schwalm, M.; Bartelle, B. B.; Xie, T.; Daher, J. I.; Singh, U. D.; Xie, K.; DiNapoli, N.; Evans, N. B.; Chung, K.; Jasanoff, A. Functional Dissection of Neural Circuitry Using a Genetic Reporter for fMRI. Nat. Neurosci. 2022, 25 (3), 390–398. 10.1038/s41593-022-01014-8.

(19) Ozbakir, H. F.; Miller, A. D. C.; Fishman, K. B.; Martins, A. F.; Kippin, T. E.; Mukherjee, A. A Protein-Based Biosensor for Detecting Calcium by Magnetic Resonance Imaging. ACS Sens. 2021, 6 (9), 3163–3169. 10.1021/acssensors.1c01085.

(20) Brustad, E. M.; Lelyveld, V. S.; Snow, C. D.; Crook, N.; Jung, S. T.; Martinez, F. M.; Scholl, T. J.; Jasanoff, A.; Arnold, F. H. Structure-Guided Directed Evolution of Highly Selective P450-Based Magnetic Resonance Imaging Sensors for Dopamine and Serotonin. J. Mol. Biol. 2012, 422 (2), 245–262. 10.1016/j.jmb.2012.05.029.

(21) Desai, M.; Sharma, J.; Slusarczyk, A. L.; Chapin, A. A.; Ohlendorf, R.; Wisniowska, A.; Sur, M.; Jasanoff, A. Hemodynamic Molecular Imaging of Tumor-Associated Enzyme Activity in the Living Brain. eLife 2021, 10, e70237. 10.7554/eLife.70237.

(22) Louie, A. Y.; Hüber, M. M.; Ahrens, E. T.; Rothbächer, U.; Moats, R.; Jacobs, R. E.; Fraser, S. E.; Meade, T. J. In Vivo Visualization of Gene Expression Using Magnetic Resonance Imaging. Nat. Biotechnol. 2000, 18 (3), 321–325. 10.1038/73780.

(23) Chalfie, M.; Tu, Y.; Euskirchen, G.; Ward, W. W.; Prasher, D. C. Green Fluorescent Protein as a Marker for Gene Expression. Science 1994, 263 (5148), 802–805. 10.1126/science.8303295.

(24) Weissleder, R.; Simonova, M.; Bogdanova, A.; Bredow, S.; Enochs, W. S.; Bogdanov, A. MR Imaging and Scintigraphy of Gene Expression through Melanin Induction. Radiology 1997, 204 (2), 425–429. 10.1148/radiology.204.2.9240530.

(25) Weissleder, R.; Moore, A.; Mahmood, U.; Bhorade, R.; Benveniste, H.; Chiocca, E. A.; Basilion, J. P. In Vivo Magnetic Resonance Imaging of Transgene Expression. Nat. Med. 2000, 6 (3), 351–354. 10.1038/73219.

(26) Faiz Kayyem, J.; Kumar, R. M.; Fraser, S. E.; Meade, T. J. Receptor-Targeted Co-Transport of DNA and Magnetic Resonance Contrast Agents. Chem. Biol. 1995, 2 (9), 615–620. 10.1016/1074-5521(95)90126-4.

(27) Mukherjee, A.; Wu, D.; Davis, H. C.; Shapiro, M. G. Non-Invasive Imaging Using Reporter Genes Altering Cellular Water Permeability. Nat. Commun. 2016, 7 (1), 13891. 10.1038/ncomms13891.

(28) Chacko, A. N.; Miller, A. D. C.; Dhanabalan, K. M.; Mukherjee, A. Exploring the Potential of Water Channels for Developing Genetically Encoded Reporters and Biosensors for Diffusion-Weighted MRI. J. Magn. Reson. 2024, 365, 107743. 10.1016/j.jmr.2024.107743.

(29) Yun, J.; Huang, Y.; Miller, A. D. C.; Chang, B. L.; Baldini, L.; Dhanabalan, K. M.; Li, E.; Li, H.; Mukherjee, A. Destabilized Reporters for Background-Subtracted, Chemically-Gated, and Multiplexed Deep-Tissue Imaging. Chem. Sci. 2024, 15 (28), 11108–11121. 10.1039/D4SC00377B.

(30) Zhang, L.; Gong, M.; Lei, S.; Cui, C.; Liu, Y.; Xiao, S.; Kang, X.; Sun, T.; Xu, Z.; Zhou, C.; Zhang, S.; Zhang, D. Targeting Visualization of Malignant Tumor Based on the Alteration of DWI Signal Generated by hTERT Promoter–Driven AQP1 Overexpression. Eur. J. Nucl. Med. Mol. Imaging 2022, 49 (7), 2310–2322. 10.1007/s00259-022-05684-1.

(31) Li, M.; Liu, Z.; Wu, Y.; Zheng, N.; Liu, X.; Cai, A.; Zheng, D.; Zhu, J.; Wu, J.; Xu, L.; Li, X.; Zhu, L.-Q.; Manyande, A.; Xu, F.; Wang, J. In Vivo Imaging of Astrocytes in the Whole Brain with Engineered AAVs and Diffusion-Weighted Magnetic Resonance Imaging. Mol. Psychiatry 2022, 1–8. 10.1038/s41380-022-01580-0.

(32) Zheng, N.; Li, M.; Wu, Y.; Kaewborisuth, C.; Li, Z.; Gui, Z.; Wu, J.; Cai, A.; Lin, K.; Su, K.-P.; Xiang, H.; Tian, X.; Manyande, A.; Xu, F.; Wang, J. A Novel Technology for in Vivo Detection of Cell Type-Specific Neural Connection with AQP1-Encoding rAAV2-Retro Vector and Metal-Free MRI. NeuroImage 2022, 258, 119402. 10.1016/j.neuroimage.2022.119402.

(33) Chowdhury, R.; Wan, J.; Gardier, R.; Rafael-Patino, J.; Thiran, J.-P.; Gibou, F.; Mukherjee, A. Molecular Imaging with Aquaporin-Based Reporter Genes: Quantitative Considerations from Monte Carlo Diffusion Simulations. ACS Synth. Biol. 2023, 12 (10), 3041–3049. 10.1021/acssynbio.3c00372.

(34) Miller, A. D. C.; Chowdhury, S. P.; Hanson, H. W.; Linderman, S. K.; Ghasemi, H. I.; Miller, W. D.; Morrissey, M. A.; Richardson, C. D.; Gardner, B. M.; Mukherjee, A. Engineering Water Exchange Is a Safe and Effective Method for Magnetic Resonance Imaging in Diverse Cell Types. J. Biol. Eng. 2024, 18 (1), 30. 10.1186/s13036-024-00424-5.

(35) Stejskal, E. O.; Tanner, J. E. Spin Diffusion Measurements: Spin Echoes in the Presence of a Time-Dependent Field Gradient. J. Chem. Phys. 2004, 42 (1), 288–292. 10.1063/1.1695690.

(36) Lasič, S.; Nilsson, M.; Lätt, J.; Ståhlberg, F.; Topgaard, D. Apparent Exchange Rate Mapping with Diffusion MRI. Magn. Reson. Med. 2011, 66 (2), 356–365. 10.1002/mrm.22782.

(37) Gao, X. J.; Chong, L. S.; Kim, M. S.; Elowitz, M. B. Programmable Protein Circuits in Living Cells. Science 2018, 361 (6408), 1252–1258. 10.1126/science.aat5062.

(38) Fink, T.; Lonzarić, J.; Praznik, A.; Plaper, T.; Merljak, E.; Leben, K.; Jerala, N.; Lebar, T.; Strmšek, Ž.; Lapenta, F.; Benčina, M.; Jerala, R. Design of Fast Proteolysis-Based Signaling and Logic Circuits in Mammalian Cells. Nat. Chem. Biol. 2019, 15 (2), 115–122. 10.1038/s41589-018-0181-6.

(39) Fernandez-Rodriguez, J.; Voigt, C. A. Post-Translational Control of Genetic Circuits Using *Potyvirus* Proteases. Nucleic Acids Res. 2016, 44 (13), 6493–6502. 10.1093/nar/gkw537.

(40) Chung, H. K.; Lin, M. Z. On the Cutting Edge: Protease-Based Methods for Sensing and Controlling Cell Biology. Nat. Methods 2020, 17 (9), 885–896. 10.1038/s41592-020-0891-z.

(41) Chen, Z.; Elowitz, M. B. Programmable Protein Circuit Design. Cell 2021, 184 (9), 2284–2301. 10.1016/j.cell.2021.03.007.

(42) Stein, V.; Alexandrov, K. Protease-Based Synthetic Sensing and Signal Amplification. Proc. Natl. Acad. Sci. 2014, 111 (45), 15934–15939. 10.1073/pnas.1405220111.

(43) Sanchez, M. I.; Ting, A. Y. Directed Evolution Improves the Catalytic Efficiency of TEV Protease. Nat. Methods 2020, 17 (2), 167–174. 10.1038/s41592-019-0665-7.

(44) Miyazaki, Y.; Imoto, H.; Chen, L.; Wandless, T. J. Destabilizing Domains Derived from the Human Estrogen Receptor. J. Am. Chem. Soc. 2012, 134 (9), 3942–3945. 10.1021/ja209933r.

(45) Banaszynski, L. A.; Chen, L.; Maynard-Smith, L. A.; Ooi, A. G. L.; Wandless, T. J. A Rapid, Reversible, and Tunable Method to Regulate Protein Function in Living Cells Using Synthetic Small Molecules. Cell 2006, 126 (5), 995–1004. 10.1016/j.cell.2006.07.025.

(46) Ha, S.-W.; Ju, D.; Xie, Y. The N-Terminal Domain of Rpn4 Serves as a Portable Ubiquitin-Independent Degron and Is Recognized by Specific 19S RP Subunits. Biochem. Biophys. Res. Commun. 2012, 419 (2), 226–231. 10.1016/j.bbrc.2012.01.152.

(47) Zhang, Q.; Schepis, A.; Huang, H.; Yang, J.; Ma, W.; Torra, J.; Zhang, S.-Q.; Yang, L.; Wu, H.; Nonell, S.; Dong, Z.; Kornberg, T. B.; Coughlin, S. R.; Shu, X. Designing a Green Fluorogenic Protease Reporter by Flipping a Beta Strand of GFP for Imaging Apoptosis in Animals. J. Am. Chem. Soc. 2019, 141 (11), 4526–4530. 10.1021/jacs.8b13042.

(48) Fasting, C.; Schalley, C. A.; Weber, M.; Seitz, O.; Hecht, S.; Koksch, B.; Dernedde, J.; Graf, C.; Knapp, E.; Haag, R. Multivalency as a Chemical Organization and Action Principle. Angew. Chem. Int. Ed. 2012, 51 (42), 10472–10498. 10.1002/anie.201201114.

(49) Mauthe, M.; Orhon, I.; Rocchi, C.; Zhou, X.; Luhr, M.; Hijlkema, K.-J.; Coppes, R. P.; Engedal, N.; Mari, M.; Reggiori, F. Chloroquine Inhibits Autophagic Flux by Decreasing Autophagosome-Lysosome Fusion. Autophagy 2018, 14 (8), 1435–1455. 10.1080/15548627.2018.1474314.

(50) Lee, D. H.; Goldberg, A. L. Proteasome Inhibitors: Valuable New Tools for Cell Biologists. Trends Cell Biol. 1998, 8 (10), 397–403. 10.1016/S0962-8924(98)01346-4.

(51) De Clercq, E.; Li, G. Approved Antiviral Drugs over the Past 50 Years. Clin. Microbiol. Rev. 2016, 29 (3), 695–747. 10.1128/CMR.00102-15.

(52) Majerová, T.; Konvalinka, J. Viral Proteases as Therapeutic Targets. Mol. Aspects Med. 2022, 88, 101159. 10.1016/j.mam.2022.101159.

(53) Rihtar, E.; Lebar, T.; Lainšček, D.; Kores, K.; Lešnik, S.; Bren, U.; Jerala, R. Chemically Inducible Split Protein Regulators for Mammalian Cells. Nat. Chem. Biol. 2023, 19 (1), 64–71. 10.1038/s41589-022-01136-x.

(54) Franko, N.; Teixeira, A. P.; Xue, S.; Charpin-El Hamri, G.; Fussenegger, M. Design of Modular Autoproteolytic Gene Switches Responsive to Anti-Coronavirus Drug Candidates. Nat. Commun. 2021, 12 (1), 6786. 10.1038/s41467-021-27072-3.

(55) Xia, S.; Lu, A. C.; Tobin, V.; Luo, K.; Moeller, L.; Shon, D. J.; Du, R.; Linton, J. M.; Sui, M.; Horns, F.; Elowitz, M. B. Synthetic Protein Circuits for Programmable Control of Mammalian Cell Death. Cell 2024, 187 (11), 2785–2800.e16. 10.1016/j.cell.2024.03.031.

(56) Rihtar, E.; Lebar, T.; Lainšček, D.; Kores, K.; Lešnik, S.; Bren, U.; Jerala, R. Chemically Inducible Split Protein Regulators for Mammalian Cells. Nat. Chem. Biol. 2023, 19 (1), 64–71. 10.1038/s41589-022-01136-x.

(57) Sun, J.; Zhang, W.; Zhao, Y.; Liu, J.; Wang, F.; Han, Y.; Jiang, M.; Li, S.; Tang, D. Conditional Control of Chimeric Antigen Receptor T-Cell Activity through a Destabilizing Domain Switch and Its Chemical Ligand. Cytotherapy 2021, 23 (12), 1085–1096. 10.1016/j.jcyt.2021.07.014.

(58) Lee, I. K.; Sharma, N.; Noguera-Ortega, E.; Liousia, M.; Baroja, M. L.; Etersque, J. M.; Pham, J.; Sarkar, S.; Carreno, B. M.; Linette, G. P.; Puré, E.; Albelda, S. M.; Sellmyer, M. A. A Genetically Encoded Protein Tag for Control and Quantitative Imaging of CAR T Cell Therapy. Mol. Ther. 2023, 31 (12), 3564–3578. 10.1016/j.ymthe.2023.10.020.

(59) Chen, J.; Lin, F.-L.; Leung, J. Y. K.; Tu, L.; Wang, J.-H.; Chuang, Y.-F.; Li, F.; Shen, H.-H.; Dusting, G. J.; Wong, V. H. Y.; Lisowski, L.; Hewitt, A. W.; Bui, B. V.; Zhong, J.; Liu, G.-S. A Drug-Tunable Flt23k Gene Therapy for Controlled Intervention in Retinal Neovascularization. Angiogenesis 2021, 24 (1), 97–110. 10.1007/s10456-020-09745-7.

(60) Maji, B.; Moore, C. L.; Zetsche, B.; Volz, S. E.; Zhang, F.; Shoulders, M. D.; Choudhary, A. Multidimensional Chemical Control of CRISPR–Cas9. Nat. Chem. Biol. 2017, 13 (1), 9–11. 10.1038/nchembio.2224.

(61) Santiago, C. P.; Keuthan, C. J.; Boye, S. L.; Boye, S. E.; Imam, A. A.; Ash, J. D. A Drug-Tunable Gene Therapy for Broad-Spectrum Protection against Retinal Degeneration. Mol. Ther. 2018, 26 (10), 2407–2417. 10.1016/j.ymthe.2018.07.016.

(62) Datta, S.; Renwick, M.; Chau, V. Q.; Zhang, F.; Nettesheim, E. R.; Lipinski, D. M.; Hulleman, J. D. A Destabilizing Domain Allows for Fast, Noninvasive, Conditional Control of Protein Abundance in the Mouse Eye – Implications for Ocular Gene Therapy. Invest. Ophthalmol. Vis. Sci. 2018, 59 (12), 4909–4920. 10.1167/iovs.18-24987.

(63) Peng, H.; Chau, V. Q.; Phetsang, W.; Sebastian, R. M.; Stone, M. R. L.; Datta, S.; Renwick, M.; Tamer, Y. T.; Toprak, E.; Koh, A. Y.; Blaskovich, M. A. T.; Hulleman, J. D. Non-Antibiotic Small-Molecule Regulation of DHFR-Based Destabilizing Domains In Vivo. Mol. Ther. Methods Clin. Dev. 2019, 15, 27–39. 10.1016/j.omtm.2019.08.002.

(64) Iwamoto, M.; Björklund, T.; Lundberg, C.; Kirik, D.; Wandless, T. J. A General Chemical Method to Regulate Protein Stability in the Mammalian Central Nervous System. Chem. Biol. 2010, 17 (9), 981–988. 10.1016/j.chembiol.2010.07.009.

(65) Nakahara, E.; Mullapudi, V.; Collier, G. E.; Joachimiak, L. A.; Hulleman, J. D. Development of a New DHFR-Based Destabilizing Domain with Enhanced Basal Turnover and Applicability in Mammalian Systems. ACS Chem. Biol. 2022, 17 (10), 2877–2889. 10.1021/acschembio.2c00518.

(66) Kwo, P. Y.; Lawitz, E. J.; McCone, J.; Schiff, E. R.; Vierling, J. M.; Pound, D.; Davis, M. N.; Galati, J. S.; Gordon, S. C.; Ravendhran, N.; Rossaro, L.; Anderson, F. H.; Jacobson, I. M.; Rubin, R.; Koury, K.; Pedicone, L. D.; Brass, C. A.; Chaudhri, E.; Albrecht, J. K. Efficacy of Boceprevir, an NS3 Protease Inhibitor, in Combination with Peginterferon Alfa-2b and Ribavirin in Treatment-Naive Patients with Genotype 1 Hepatitis C Infection (SPRINT-1): An Open-Label, Randomised, Multicentre Phase 2 Trial. The Lancet 2010, 376 (9742), 705–716. 10.1016/S0140-6736(10)60934-8.

(67) McHutchison, J. G.; Manns, M. P.; Muir, A. J.; Terrault, N. A.; Jacobson, I. M.; Afdhal, N. H.; Heathcote, E. J.; Zeuzem, S.; Reesink, H. W.; Garg, J.; Bsharat, M.; George, S.; Kauffman, R. S.; Adda, N.; Di Bisceglie, A. M. Telaprevir for Previously Treated Chronic HCV Infection. N. Engl. J. Med. 2010, 362 (14), 1292–1303. 10.1056/NEJMoa0908014.

(68) Wehr, M. C.; Laage, R.; Bolz, U.; Fischer, T. M.; Grünewald, S.; Scheek, S.; Bach, A.; Nave, K.-A.; Rossner, M. J. Monitoring Regulated Protein-Protein Interactions Using Split TEV. Nat. Methods 2006, 3 (12), 985–993. 10.1038/nmeth967.

(69) Gradišar, H.; Jerala, R. *De Novo* Design of Orthogonal Peptide Pairs Forming Parallel Coiled-coil Heterodimers. J. Pept. Sci. 2011, 17 (2), 100–106. 10.1002/psc.1331.

(70) Erofeev, A. I.; Vinokurov, E. K.; Vlasova, O. L.; Bezprozvanny, I. B. GCaMP, a Family of Single-Fluorophore Genetically Encoded Calcium Indicators. J. Evol. Biochem. Physiol. 2023, 59 (4), 1195–1214. 10.1134/S0022093023040142.

(71) Stornaiuolo, M.; Lotti, L. V.; Borgese, N.; Torrisi, M.-R.; Mottola, G.; Martire, G.; Bonatti, S. KDEL and KKXX Retrieval Signals Appended to the Same Reporter Protein Determine Different Trafficking between Endoplasmic Reticulum, Intermediate Compartment, and Golgi Complex. Mol. Biol. Cell 2003, 14 (3), 889–902. 10.1091/mbc.e02-08-0468.

(72) Praznik, A.; Fink, T.; Franko, N.; Lonzarić, J.; Benčina, M.; Jerala, N.; Plaper, T.; Roškar, S.; Jerala, R. Regulation of Protein Secretion through Chemical Regulation of Endoplasmic Reticulum Retention Signal Cleavage. Nat. Commun. 2022, 13 (1), 1323. 10.1038/s41467-022-28971-9.

(73) Wang, X.; Kang, L.; Kong, D.; Wu, X.; Zhou, Y.; Yu, G.; Dai, D.; Ye, H. A Programmable Protease-Based Protein Secretion Platform for Therapeutic Applications. Nat. Chem. Biol. 2024, 20 (4), 432–442. 10.1038/s41589-023-01433-z.

(74) Vlahos, A. E.; Kang, J.; Aldrete, C. A.; Zhu, R.; Chong, L. S.; Elowitz, M. B.; Gao, X. J. Protease-Controlled Secretion and Display of Intercellular Signals. Nat. Commun. 2022, 13 (1), 912. 10.1038/s41467-022-28623-y.

(75) Gray, D. C.; Mahrus, S.; Wells, J. A. Activation of Specific Apoptotic Caspases with an Engineered Small-Molecule-Activated Protease. Cell 2010, 142 (4), 637–646. 10.1016/j.cell.2010.07.014.

(76) Wehr, M. C.; Reinecke, L.; Botvinnik, A.; Rossner, M. J. Analysis of Transient Phosphorylation-Dependent Protein-Protein Interactions in Living Mammalian Cells Using Split-TEV. BMC Biotechnol. 2008, 8 (1), 55. 10.1186/1472-6750-8-55.

(77) Wehr, M. C.; Rossner, M. J. Split Protein Biosensor Assays in Molecular Pharmacological Studies. Drug Discov. Today 2016, 21 (3), 415–429. 10.1016/j.drudis.2015.11.004.

(78) Stanton, B. Z.; Chory, E. J.; Crabtree, G. R. Chemically Induced Proximity in Biology and Medicine. Science 2018, 359 (6380), eaao5902. 10.1126/science.aao5902.

(79) Barnea, G.; Strapps, W.; Herrada, G.; Berman, Y.; Ong, J.; Kloss, B.; Axel, R.; Lee, K. J. The Genetic Design of Signaling Cascades to Record Receptor Activation. Proc. Natl. Acad. Sci. 2008, 105 (1), 64–69. 10.1073/pnas.0710487105.

(80) Kroeze, W. K.; Sassano, M. F.; Huang, X.-P.; Lansu, K.; McCorvy, J. D.; Giguère, P. M.; Sciaky, N.; Roth, B. L. PRESTO-Tango as an Open-Source Resource for Interrogation of the Druggable Human GPCRome. Nat. Struct. Mol. Biol. 2015, 22 (5), 362–369. 10.1038/nsmb.3014.

(81) Wintgens, J. P.; Wichert, S. P.; Popovic, L.; Rossner, M. J.; Wehr, M. C. Monitoring Activities of Receptor Tyrosine Kinases Using a Universal Adapter in Genetically Encoded Split TEV Assays. Cell. Mol. Life Sci. CMLS 2019, 76 (6), 1185–1199. 10.1007/s00018-018-03003-2.

(82) Djannatian, M. S.; Galinski, S.; Fischer, T. M.; Rossner, M. J. Studying G Protein-Coupled Receptor Activation Using Split-Tobacco Etch Virus Assays. Anal. Biochem. 2011, 412 (2), 141–152. 10.1016/j.ab.2011.01.042.

(83) Huang, H.; Zhang, X.; Lv, J.; Yang, H.; Wang, X.; Ma, S.; Shao, R.; Peng, X.; Lin, Y.; Rong, Z. Cell-Cell Contact-Induced Gene Editing/Activation in Mammalian Cells Using a synNotch-CRISPR/Cas9 System. Protein Cell 2020, 11 (4), 299–303. 10.1007/s13238-020-00690-1.

(84) Gao, X. J.; Chong, L. S.; Kim, M. S.; Elowitz, M. B. Programmable Protein Circuits in Living Cells. Science 2018, 361 (6408), 1252–1258. 10.1126/science.aat5062.

(85) Aldrete, C. A.; An, C.; Call, C. C.; Gao, X. J.; Vlahos, A. E. Perspectives on Synthetic Protein Circuits in Mammalian Cells. Curr. Opin. Biomed. Eng. 2024, 32, 100555. 10.1016/j.cobme.2024.100555.

(86) Cui, M.; Lan, T.-H.; Zhou, Y. Harnessing Viral Proteases for Cellular and Molecular Engineering. Chemistry–Methods n/a (n/a), 2400098. 10.1002/cmtd.202400098.

(87) Rohner, E.; Yang, R.; Foo, K. S.; Goedel, A.; Chien, K. R. Unlocking the Promise of mRNA Therapeutics. Nat. Biotechnol. 2022, 40 (11), 1586–1600. 10.1038/s41587-022-01491-z.

(88) Patrick, P. S.; Hammersley, J.; Loizou, L.; Kettunen, M. I.; Rodrigues, T. B.; Hu, D.-E.; Tee, S.-S.; Hesketh, R.; Lyons, S. K.; Soloviev, D.; Lewis, D. Y.; Aime, S.; Fulton, S. M.; Brindle, K. M. Dual-Modality Gene Reporter for in Vivo Imaging. Proc. Natl. Acad. Sci. 2014, 111 (1), 415–420. 10.1073/pnas.1319000111.

(89) Kelly, J. J.; Saee-Marand, M.; Nyström, N. N.; Evans, M. M.; Chen, Y.; Martinez, F. M.; Hamilton, A. M.; Ronald, J. A. Safe Harbor-Targeted CRISPR-Cas9 Homology-Independent Targeted Integration for Multimodality Reporter Gene-Based Cell Tracking. Sci. Adv. 2021, 7 (4), eabc3791. 10.1126/sciadv.abc3791.

(90) Cohen, B.; Dafni, H.; Meir, G.; Harmelin, A.; Neeman, M. Ferritin as an Endogenous MRI Reporter for Noninvasive Imaging of Gene Expression in C6 Glioma Tumors. Neoplasia 2005, 7 (2), 109–117. 10.1593/neo.04436.

(91) Perlman, O.; Ito, H.; Gilad, A. A.; McMahon, M. T.; Chiocca, E. A.; Nakashima, H.; Farrar, C. T. Redesigned Reporter Gene for Improved Proton Exchange-Based Molecular MRI Contrast. Sci. Rep. 2020, 10, 20664. 10.1038/s41598-020-77576-z.

(92) Ohlendorf, R.; Wiśniowska, A.; Desai, M.; Barandov, A.; Slusarczyk, A. L.; Li, N.; Jasanoff, A. Target-Responsive Vasoactive Probes for Ultrasensitive Molecular Imaging. Nat. Commun. 2020, 11 (1), 2399. 10.1038/s41467-020-16118-7.

(93) Bourdeau, R. W.; Lee-Gosselin, A.; Lakshmanan, A.; Farhadi, A.; Kumar, S. R.; Nety, S. P.; Shapiro, M. G. Acoustic Reporter Genes for Noninvasive Imaging of Microorganisms in Mammalian Hosts. Nature 2018, 553 (7686), 86–90. 10.1038/nature25021.

(94) Lakshmanan, A.; Jin, Z.; Nety, S. P.; Sawyer, D. P.; Lee-Gosselin, A.; Malounda, D.; Swift, M. B.; Maresca, D.; Shapiro, M. G. Acoustic Biosensors for Ultrasound Imaging of Enzyme Activity. Nat. Chem. Biol. 2020, 16 (9), 988–996. 10.1038/s41589-020-0591-0.

(95) Farhadi, A.; Sigmund, F.; Westmeyer, G. G.; Shapiro, M. G. Genetically Encodable Materials for Non-Invasive Biological Imaging. Nat. Mater. 2021, 20 (5), 585–592. 10.1038/s41563-020-00883-3.

