## Supporting Information for "A programmable genetic platform for engineering noninvasive biosensors"

Affiliations:

### Materials

Reagents for polymerase chain reaction (PCR) and Gibson multi-gene assembly were procured from New England Biolabs. Doxycycline hyclate, penicillin/streptomycin ($10^4$ units/mL penicillin and 10 mg/mL streptomycin), β-mercaptoethanol, MG132, chloroquine diphosphate, trimethoprim, leupeptin, boceprevir, and telaprevir were obtained from Millipore Sigma. Shield-1 was purchased from Aobious Inc. Ionomycin was purchased from Stem Cell Technologies. Polyethyleneimine (linear, 25 kDa) was obtained from Polysciences. The Lenti-X viral concentrator was sourced from Takara Bio. RPMI 1640 cell culture medium, Ham's F-12K (Kaighn's) medium, and DMEM containing high glucose (4.5 g/L), L-glutamine, and sodium pyruvate, was acquired from Thermo Fisher Scientific. Polybrene (hexadimethrine bromide) and RIPA lysis buffer were purchased from Santa Cruz Biotechnology. Fetal bovine serum (FBS), Pierce BCA protein assay kit, and 4′,6-diamidino-2-phenylindole lactate salt (DAPI) were procured from Thermo Fisher Scientific. Reagents for denaturing gel electrophoresis, including pre-cast gels, Laemmli buffer, Tris-buffered saline with Tween-20 (TBS-T), non-fat dried milk, and Clarity Western ECL reagents, were obtained from Bio-Rad. The WesternSure chemiluminescent pre-stained protein ladder for immunoblotting was acquired from LI-COR Biosciences. Mem-PER Plus membrane protein extraction kit was obtained from Thermo Fisher Scientific. 10x PBS was obtained from Apex Bioresearch Products. PBST was made by adding 0.1% (*w*/*v*) Tween® 20 purchased from Thermofisher Scientific to 1x PBS. Amicon Ultra-15 centrifugal filters (10 kDa MWCO) were purchased from Millipore Sigma. Mouse α-FLAG primary antibody was procured from Sigma-Aldrich (#F1804). Horseradish peroxidase (HRP)-conjugated (#TL280988) and Alexa Fluor 647 (#A21235)-conjugated goat α-mouse IgG secondary antibodies were purchased from Thermo Fisher Scientific. Rabbit α-Na/K ATPase primary antibody (#AB76020) and HRP-conjugated goat anti-Rabbit IgG secondary antibody (#AB205718) were purchased from Abcam.

### Molecular biology

All plasmids generated in this work are described in Table 1. Sequences of the main genetic components used to construct and test AURCAS are listed in Table 2. Plasmids harboring the ER50 and FKBP12-based destabilizing domains (Addgene #236279) have been described in a previous publication[1] from our lab. *E. coli* DHFR was obtained from Addgene (#29326) and the following mutations were introduced to convert it into a destabilizing domain as described before[2]: H12R, T18N, V19A, S67G, W74R, T113S, E120D, and Q146L. Plasmid containing the FlipGFP TEVP sensor was obtained from Addgene (#124429). FKBP and Frb split TEV systems were also obtained from Addgene(#137831, #137832). All the other genes were synthesized by Integrated DNA Technologies (IDT). The genes of interest were amplified using Q5® High-Fidelity DNA Polymerase and cloned into a lentiviral transfer plasmid under the control of either a constitutive promoter, EF1α (Addgene #60058) or a doxycycline-inducible minimal CMV promoter (Addgene #26431). Constitutively expressed fluorescent reporters, including EBFP, EGFP, dTomato or mCherry, were included in our lentiviral vectors to allow sorting for successfully transduced cells via fluorescence activated cell sorting (FACS). A FLAG tag was added to an extracellular loop in hAQP1 (immediately after Q43) to facilitate the detection of surface-express hAqp1 using α-FLAG antibodies. All final plasmids were assembled via Gibson assembly and verified by whole-plasmid nanopore sequencing (Plasmidsaurus). PCR-based amplification and Gibson assembly were conducted as described in our earlier works[1,3].

**Cell Culture, Transfections, and Transductions**

Cell lines were obtained from American Type Cell Culture Collection (U87, Jurkat, PC-12, MDA-MB-231, HEK 293T) or from Clontech (CHO TetON). CHO Tet ON, 293T, and U87 cells were grown in DMEM supplemented with 10% FBS, 100 unit/mL penicillin, and 100 µg/mL streptomycin. Jurkat cells were grown in RPMI supplemented with 10% FBS, and 100 unit/mL penicillin and 100 µg/mL streptomycin. PC-12 cells were grown in F-12K medium supplemented with FBS to a final concentration of 2.5%, horse serum to a final concentration of 15%, 100 unit/mL penicillin, and 100 µg/mL streptomycin. All cells were routinely cultured in a 37 °C humidified incubator containing 5% $CO_2$.

HEK 293T cells were used for packaging the lentivirus as described in our earlier works[1,3]. Briefly, cells were grown to 50-60% confluency in a 10 cm plate and transfected with the following three plasmids: 22 µg of packaging plasmid expressing the capsid genes from a CMV promoter, 22 µg of transfer plasmid harboring the gene of interest flanked by long terminal repeat sequences, and 4.5 µg of VSV-G plasmid that confers broad tropism to the lentiviral particles. Transfection were performed with 25 kDa linear polyethyleneimine at a concentration of 2.58 mg PEI per mg DNA. After 24 hours of transfection, the transfection media was supplemented with 100 uL of sodium butyrate to yield a final concentration of 10 mM. Virus production was allowed to continue for 48 h, following which the media was centrifuged at 500 x g for 10 min to remove residual cells and debris and concentrated by mixing with one-third volume of Lenti-X at 4 °C for 24 h. The viral particles were separated by centrifuging the solution at 1500 x g for 45 min at 4 °C and resuspending the pellet in 0.2 mL sterile phosphate buffered saline (PBS).

For stable transductions, cells were grown in 6-well plates to 70-80% confluency. Spent media was aspirated and replaced with 1 mL of fresh media containing 30 µL aliquots of the virus together with 8 ug/mL polybrene. Cells were spinfected at 1050 x g for 90 min at 30 °C, cultured at 37 °C for 48 h, before finally transferring them to a 10 cm plate and growing till confluency. The confluent cells were harvested for sorting stably transduced cells using FACS.

**Fluorescence activated cell sorting**

Cells grown to confluence in 10 cm dishes were harvested, pelleted at 300 × g for 5 min, resuspended in 2 mL of media, and filtered through a 40 µm pluriStrainer Mini cell strainer. Sorting was performed on a Sony MA900 cell sorter using a 100 µm chip. The instrument was calibrated using the manufacturer's automated protocol using Sony calibration beads. Gates were defined using negative control cells based on FSC vs SSC (cell size and granularity), FSC-H vs FSC-W (singlet discrimination), and FSC-H vs fluorescence (EBFP, EGFP, mCherry or dTomato). The emission was collected using a 450/50nm band-width filter for EBFP, 525/50nm for EGFP, and 617/30nm for mCherry or dTomato. Fluorescent populations comprising between 20,000 to 100,000 events were sorted into tubes containing at least 5 mL media. Fluorescent populations were gated to include only those exhibiting at least a 10-fold increase in fluorescence intensity (log scale) over the baseline population. Samples failing to meet this threshold were excluded and remade. In case of multiple stable transductions, the sensor(always on EGFP) was transduced last, and the cells were sorted after each transduction. A representative gating scheme involving three transductions is depicted in Figure S2. After sorting, cells were centrifuged (300 × g, 10 min), media was reduced to ~100 µL, and cells were resuspended in 2 mL (sorted) media and grown to confluency in a 6 well plate, before moving it to a 10 cm plate.

**Drug treatment**

Stock solution of doxycycline hyclate was prepared at 10 mg/mL in sterile deionized water. Stock solution of shield-1 was prepared at 1 mM in DMSO. MG132, Trimethoprim, boceprevir, and telaprevir were prepared in DMSO at 10 mM. Rapamycin was prepared in DMSO at 2.5 mM. 100mM chloroquine diphosphate and 200mM Leupeptin stocks were prepared in water. Prior to MRI, cells were seeded to reach 70-80% confluency in 48 h. Small-molecule drugs were added 24 h prior to imaging the cells at the following working concentrations: shield-1 (1µM), rapamycin (2.5µM), boceprevir (10µM), telaprevir (10µM), doxycycline (5µg/mL), and ionomycin (0.5 µM). Degradation inhibitors including MG132 (10 µM), chloroquine diphosphate (100 µM) and Leupeptin (200µM) were added 16 h prior to imaging using MRI. Vehicle-treated cells were used as negative controls to estimate non-specific changes in diffusivity.

**Western Blotting**

To prepare whole-cell extracts, cells were lysed using RIPA buffer. Membrane extracts were prepared using the Mem-PER Plus membrane protein extraction kit in accordance with the manufacturer's protocol. Cell extracts were concentrated utilizing a 10 kDa MWCO filter. The total protein concentration was quantified using the BCA assay, and approximately 10 µg of protein was denatured in Laemmli buffer supplemented with 5% β-mercaptoethanol by sonicating for 10 min at room temperature in a benchtop ultrasonic bath. Notably, thermal denaturation was avoided as heating induced protein aggregation, thereby preventing the samples from entering the gel. The denatured samples were resolved by gel electrophoresis at 120 V for 50 min and subsequently transferred to a nitrocellulose membrane using the Bio-Rad Trans-Blot Turbo dry transfer system. Membranes were blocked for 1 h by incubation in blocking solution (5% nonfat dried milk in TBS-T) at room temperature and subsequently incubated with primary antibodies (1 µg/mL mouse anti-FLAG or 0.05 µg/mL rabbit anti-Na+/K+-ATPase) in blocking solution overnight at 4°C. The membrane was then washed three times in TBS-T and incubated with the secondary antibody (0.5 µg/mL HRP-conjugated goat anti-mouse or anti-rabbit IgG) in blocking solution at room temperature for another 2 h. Finally, the membranes were washed three times in TBS-T and imaged using Clarity Western ECL reagents using an iBright FL1500 imaging system (Thermo Fisher Scientific).

**Immunofluorescence imaging**

CHO cells were seeded to reach 40-50% confluency in 35 mm glass bottom dish containing a 10 mm #1 (130-160 µm thickness) cover glass micro-well insert (Greiner Bio-One Cellview 35/10 mm culture dish, #627860). After reaching confluency, the cells were fixed by replacing the growth medium with 1 mL 4% paraformaldehyde and incubating for 15 min at room temperature in a fume hood. Each well of cells was washed three times with 2 mL PBS-T for 5 min and blocked for 1 h by incubating in blocking solution (2% BSA, 5% goat serum in PBS-T) with mild shaking. Subsequently, the blocking solution was replaced with 1 mL primary antibody solution (mouse anti-FLAG diluted to 0.5 ug/ml in blocking solution) and incubated at 4 °C overnight in an orbital shaker. Each well of cells was washed thrice using 2mL PBS-T (5 min per wash) and then incubated with 1 mL secondary antibody (goat α-mouse IgG H&L conjugated with Alexa Fluor® 647) diluted to 1 ug/ml in 2% BSA in PBS-T. Each plate was wrapped in aluminum foil and shaken at room temperature for 2 h in an orbital shaker. Cells were washed twice using 2 mL PBS-T (5 min per wash). Next, aqueous solution of DAPI lactate (5mg/ml) was diluted 1:2000 in PBS-T and 1 mL of the diluted solution was added to each well, incubated for 5 min, and removed by washing twice with PBS-T (3 min per wash). Lastly, 1 mL of PBS-T was added to each well before imaging. Confocal microscopy was performed using a Leica Dmi8 SP8 resonant scanning microscope equipped with an HC PL APO 63x/1.40 oil-immersion objective using Type F immersion oil. A 405 nm laser line was used for DAPI excitation, and the emitted light was detected between 410 nm and 483 nm. A 653 nm laser line was used for Alexa Fluor 647 excitation, and the emitted light was detected between 665 nm and 779 nm.

**Quantitative Reverse Transcription PCR (RT-qPCR)**

AQP1 and TEV expressing cells were seeded in six-well plates and grown to 80% confluency prior to isolating RNA using Zymo Direct-Zol RNA miniprep kit following the manufacturer protocol. One microgram of RNA was reverse-transcribed using iScript™ cDNA Synthesis Kit, following the supplied protocol. The resulting cDNA was diluted 1:10 in nuclease-free water, and gene-specific quantitative PCR (RT-qPCR) was performed with PowerUp™ SYBR™ Green Master Mix using 20 ng of cDNA per reaction on a CFX96 Touch Real-Time PCR Detection System (Bio-Rad). Cq values were obtained by regression fitting in CFX Maestro Software (Bio-Rad). Primers were designed in Primer3, with the forward primer (5′-CTCCTGGCTATTGACTACACTGG-3′) and the reverse primer(5′-GATGAAGTCGTAGATGAGTACAGCC-3′) annealing to the AQP transgene. β-Actin primers used for normalization were 5′-CCCCATTGAACACGGCATTG-3′ (forward) and 5′-AGGTCTCAAACATGATCTGGGT-3′ (reverse). Primer melting temperatures were optimized to 60 °C. Amplification specificity was confirmed by melt-curve analysis and by agarose gel electrophoresis of the amplicon. Aqp1 transcript abundance was calculated using the $2^{-\Delta\Delta Cq}$ method[4] relative to TEV-only controls, with β-actin as the internal reference gene.

**Live-cell diffusion-weighted imaging**

The protocol for live-cell diffusion-weighted imaging has been described in detail in our earlier works[3,5,6]. Briefly, in preparation for MRI, adherent cells were trypsinized, resuspended in DMEM, centrifuged at 350 x g, and resuspended again in 200 uL sterile PBS. The cells were transferred into 0.2 mL tubes and spun down 500 x g to form live-cell pellets, which were used for diffusion-weighted imaging. The tubes with cell pellets were placed in agarose (1% w/v) molds in a 3D-printed MRI phantom. The pellets were imaged using a 66 mm diameter coil in a Bruker 7T vertical-bore MRI scanner. Stimulated echo diffusion-weighted images of cell pellets were obtained in the axial plane using echo time, $T_E$ = 18 ms; repetition time, $T_R$ = 1000 ms; gradient duration; $\delta$ = 5 ms, gradient separation, $\Delta$ = 300 ms; matrix size = 128 x 128; field of view (FOV) = 5.08 x 5.08 $cm^2$; slice thickness = 1-2 mm; and number of averages = 5. Diffusion-sensitizing gradients were applied in the readout direction (dorsal-ventral axis) in the plane orthogonal to the axial slice-selection gradient using nominal b-values of 0-0.8 $ms/\mu m^2$, which correspond to effective b-values (viz., b-values corrected to include the contribution of imaging gradients) in the range of 1-3 $ms/\mu m^2$. Diffusion-weighted intensity for a given b-value was estimated by computing the average intensity of all voxels inside a region of interest (ROI) within the axial cross section of a cell pellet. The slope of the logarithmic decay in mean signal intensity as a function of b-value was used to calculate the apparent diffusivity (D). To generate a diffusion map, apparent diffusivity was computed for each voxel in an ROI. Least-squares regression fitting was performed using the *fitnlm* function in Matlab (R2022b).

**Software**

Python or Matlab were used for least-squares regression fitting to estimate diffusivity and generate live-cell diffusion maps. Python and Microsoft Excel were used for statistical analysis, including bootstrapping and hypothesis testing. To compare sample means between two groups, a two-sided Student's t-test was used to calculate P-values. When comparing sample across more than two groups, a one-way analysis of variance (ANOVA) followed by Tukey's HSD test was applied. The level of statistical significance was set at $P < 0.05$. Python was used for data visualization (bar graphs and scatter plots). Leica Application Suite X (LAS X) and ParaVision 6.0 were used to respectively acquire raw images from confocal imaging and MRI. All additional image analysis, such as selecting appropriate regions of interest (ROI) and fields-of-view encompassing single cells (microscopy) or cell pellets (MRI), quantifying pixel intensities, pseudo-coloring, and smoothing, was performed using Fiji. Inkscape was employed to collate and prepare the final figures, while Biorender was used to create graphic representations of the proteasome, lysosome, and the design-build-test workflow as depicted in Figure 1.

### Supplementary Figures

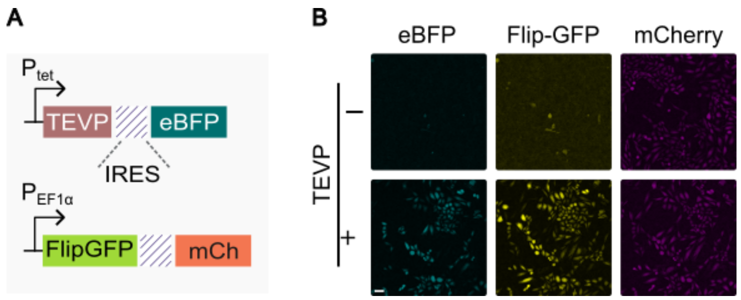


**Figure S1. Validation of TEVP activity**.

(A) Genetic constructs encoding TEVP and FlipGFP, with the latter functioning as a sensor for TEVP activity.

(B) Representative fluorescence images illustrating the activation of the FlipGFP sensor in cells expressing TEVP. Scale bar is 20 μm

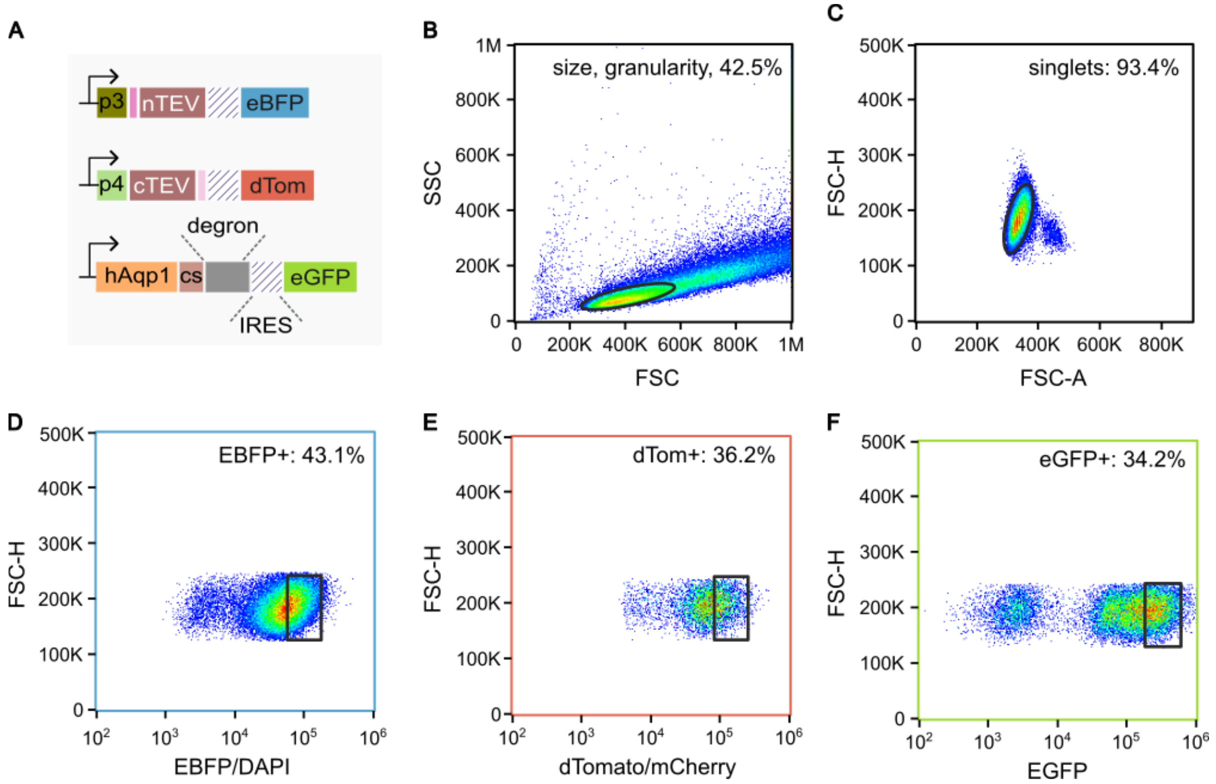


**Figure S2. Representative fluorescence-activated cell sorting of transduced cell lines.**

(A) Genetic constructs encoding the 3-color split TEV-based URCAS circuit for detecting interactions between p3 and p4 coiled coils.

(B) – (F) illustrate the three-color flow sorting strategy for cells transduced with the constructs in (A), along with the corresponding sorting gates. Fluorescence emission from eBFP, eGFP, and dTomato was detected using the following filters: eBFP/DAPI, 450/50 nm bandwidth; eGFP, 525/50 nm bandwidth; and mCherry, 617/30 nm bandwidth.

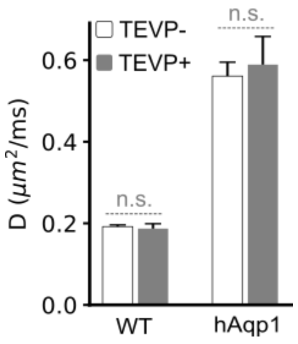


**Figure S3. Effect of TEVP expression on cellular diffusivity.**

Diffusivities of CHO cells engineered to induce TEVP expression, either in the absence of hAqp1 or with hAqp1 expressed without a fused destabilizing domain (DD). Error bars represent standard deviation from $n \geq 3$ measurements; n.s. indicates $P \geq 0.05$

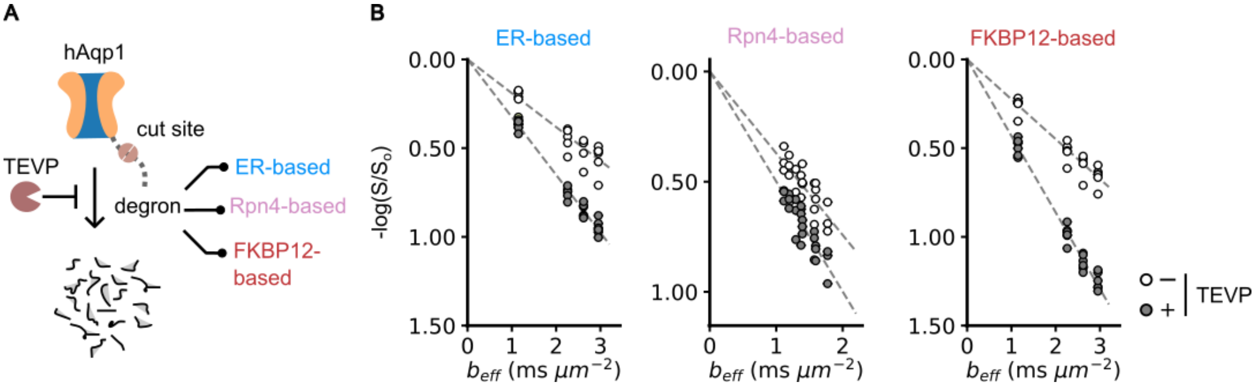


**Figure S4. Evaluation of hAqp1-DD fusions for optimal TEVP response.**

(A) Schematic representation of the mechanism by which hAqp1-based MRI signals, quantified in terms of diffusivity (D, μm$^2$/ms), are activated via proteolytic cleavage of a destabilizing domain (DD).

(B) Representative plots depicting the logarithmic decay in normalized diffusion-weighted signal intensity, both with and without the induction of TEVP expression, as a function of the diffusion-weighting factor ($b_{eff}$) in CHO cells engineered to express TEVP and various hAqp1-DD fusions. The rate of diffusion (diffusivity) was determined from the slope of the decay curve.

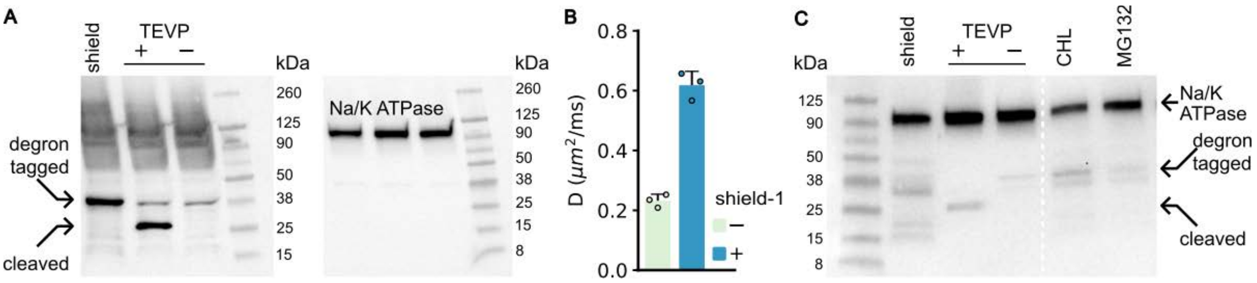


**Figure S5. Biochemical assays of TEVP-mediated modulation of MRI signals in the sensor.**

(A) Representative Western blot of membrane extracts from CHO cells engineered to express hAqp1 fused at its C-terminus to the TEVP-cleavable FKBP12-DD, with and without TEVP induction. Membrane lysates from cells treated with shield-1, which stabilizes the FKBP12-DD, are included as a positive control. The blot was initially incubated with antibodies against Na+/K+ ATPase, imaged, stripped, and subsequently incubated with an α-FLAG antibody to visualize hAqp1 expression.

(B) Quantification of diffusivities in CHO cells expressing hAqp1 fused at its C-terminus to FKBP12-DD with and without shield-1 incubation for 24 h. Error bars represent standard deviation from n ≥ 3 measurements; *** denotes P < 0.001.

(C) Representative Western blot of whole-cell lysates prepared from cells expressing DD-URCAS, with or without TEVP expression, and in the presence of the lysosomal inhibitor, chloroquine or the proteasomal inhibitor, MG132.

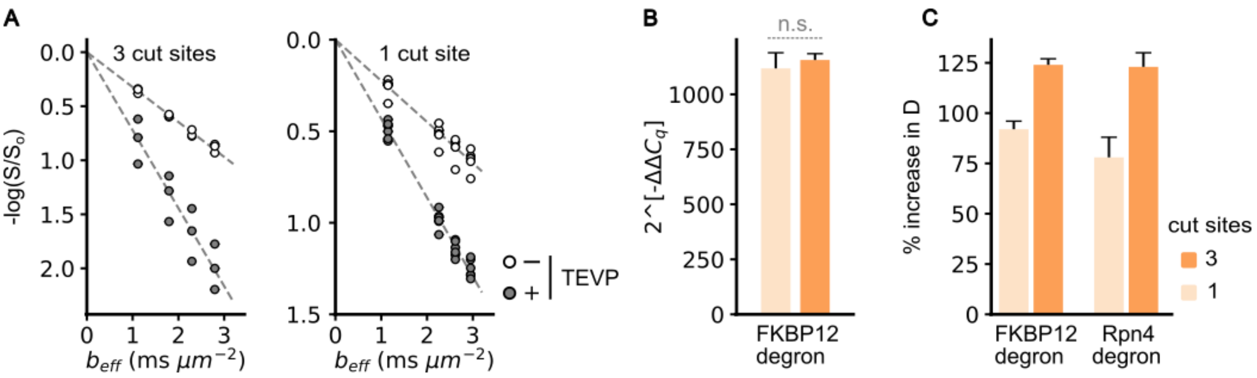


**Figure S6. Modulation of TEVP-induced change in diffusivity by varying the number of protease cleavage sites**.

(A) Representative plots depicting the logarithmic decay in normalized diffusion-weighted signal intensity, both with and without the induction of TEVP expression, as a function of the diffusion-weighting factor ($b_{eff}$) in CHO cells engineered to express TEVP and hAqp1 fused to FKBP12-DD separated by two or three cleavage sits for TEVP. The rate of diffusion (diffusivity) was determined from the slope of the decay curve.

(B) Quantification of hAqp1 expression (via qRT-PCR) in CHO cells expressing hAqp1 fused to FKBP12-DD, separated by two or three cleavage sites for TEVP. Actin served as the housekeeping gene, and relative expression was quantified using the $\Delta\Delta C_q$ method.

(C) Quantification of the percentage increase in diffusivities upon TEVP expression in CHO cells expressing hAqp1 attached to FKBP12 or Rpn4 DDs, separated by one or three cut sites.

Error bars represent standard deviation from n ≥ 3 measurements. Statistical significance is denoted by *, which indicates $P < 0.05$; ** denoting $P < 0.01$; *** denoting $P < 0.001$; while n.s. indicates $P \geq 0.05$.

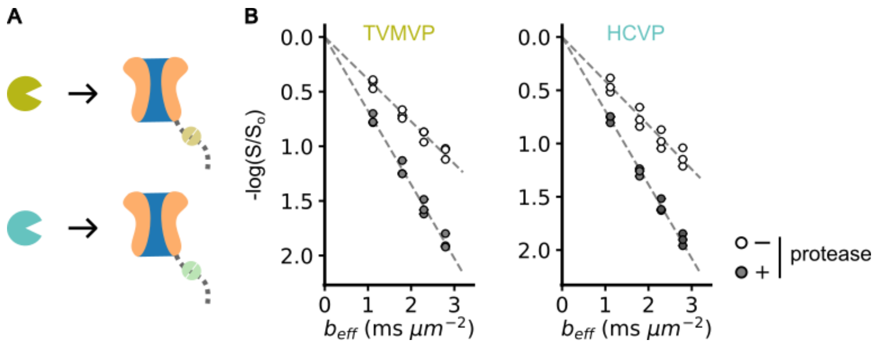


**Figure S7. Modularity of the DD-URCAS**.

(A) Schematic illustrating the adaptation of DD-URCAS to detect distinct proteases.

(B) Representative plots depicting the logarithmic decay in normalized diffusion-weighted signal intensity, both with and without the induction of TEVP expression, as a function of the diffusion-weighting factor ($b_{eff}$) in CHO cells engineered to express TVMVP- and HCVP-responsive DD-URCAS. The rate of diffusion (diffusivity) was determined from the slope of the decay curve.

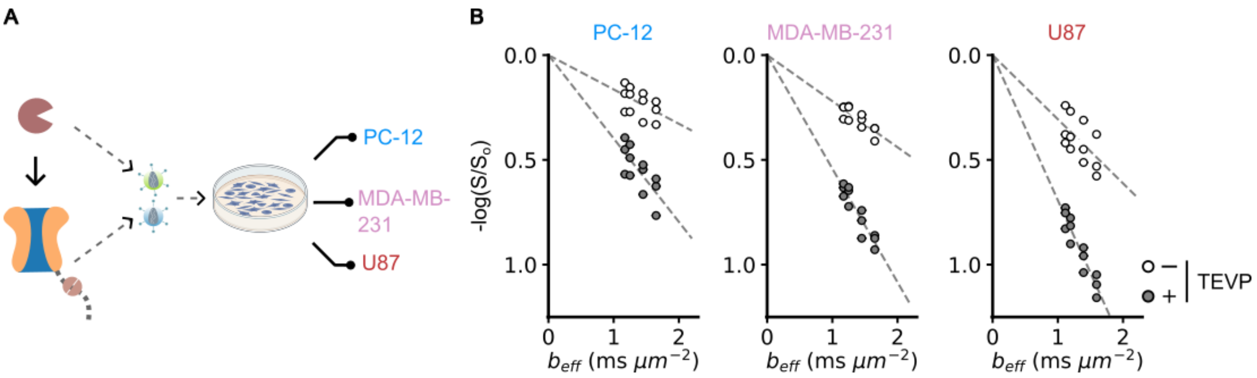


**Figure S8. Versatility of DD-URCAS across diverse cell types**.

(A) Schematic illustrating the application of DD-URCAS in distinct cell types.

(B) Representative plots depicting the logarithmic decay in normalized diffusion-weighted signal intensity, both with and without the induction of TEVP expression, as a function of the diffusion-weighting factor ($b_{eff}$) in different cell types engineered to express TVP-responsive DD-URCAS. The rate of diffusion (diffusivity) was determined from the slope of the decay curve.

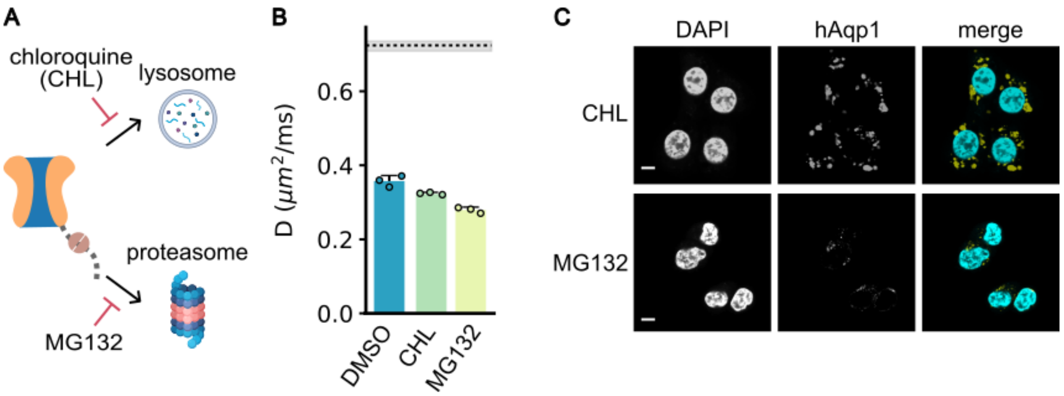


**Figure S9. Mechanistic Analysis of the degradation pathway of DD-URCAS**.

(A) Schematic illustrating the putative degradation pathways of DD-URCAS, and their inhibition by small-molecule drugs, chloroquine, and MG132

(B) Quantification of diffusivities in CHO cells expressing DD-URCAS after incubating with 0.1% DMSO (vehicle), 100 µM chloroquine, or 10µM MG132 for 24 h. The shaded band represents the 95% confidence interval of the mean diffusivity measured in DD-URCAS expressing CHO cells in the TEVP-on state.

(C) Representative confocal images of CHO cells expressing DD-URCAS after incubation with chloroquine, or MG132. Scale bar is 10µm.

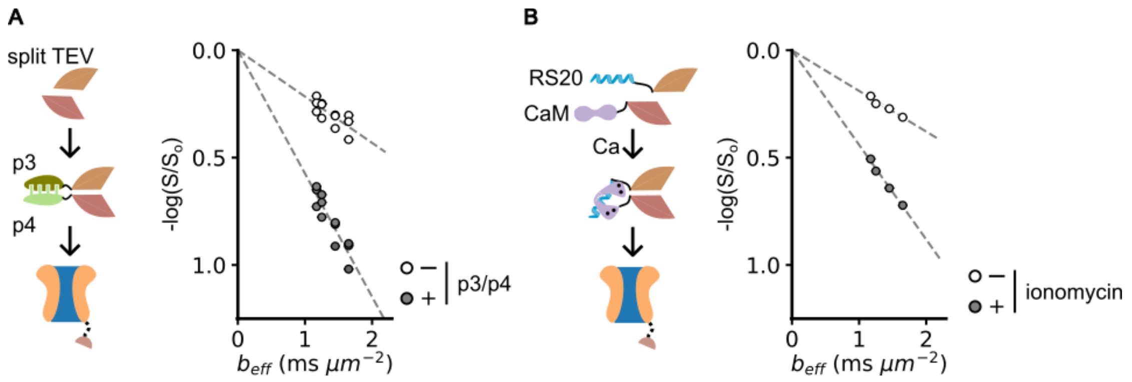


**Figure S10. Programmable biosensor design based on DD-URCAS**.

(A) Schematic illustrating the adaptation of DD-URCAS to detect biomolecular binding via reconstitution of split TEVP activity.

(B) Representative plots depicting the logarithmic decay in normalized diffusion-weighted signal intensity as a function of the diffusion-weighting factor ($b_{eff}$) in CHO cells engineered to express split TEVP-based DD-URCAS heterodimerization sensor. The rate of diffusion (diffusivity) was determined from the slope of the decay curve.

(C) Schematic illustrating the adaptation of DD-URCAS to detect intracellular calcium.

(D) Representative plots depicting the logarithmic decay in normalized diffusion-weighted signal intensity as a function of the diffusion-weighting factor ($b_{eff}$) in CHO cells engineered to express split TEVP-based calcium sensor. The rate of diffusion (diffusivity) is determined by the slope of the decay.

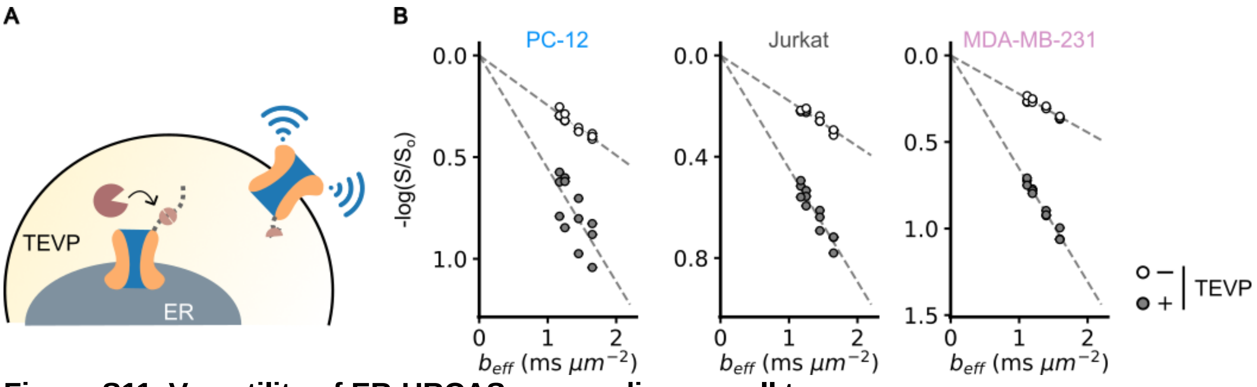


**Figure S11. Versatility of ER-URCAS across diverse cell types**

(A) Schematic illustrating the mechanism by which hAqp1 based MRI signals are activated through proteolytic cleavage of an ER retention tag (KKYL).

(B) Representative plots depicting the logarithmic decay in normalized diffusion-weighted signal intensity as a function of the diffusion-weighting factor (*$b_{eff}$*) in three different cell types expressing ER-URCAS with or without TEVP induction. The rate of diffusion (diffusivity) was determined from the slope of the decay curve.

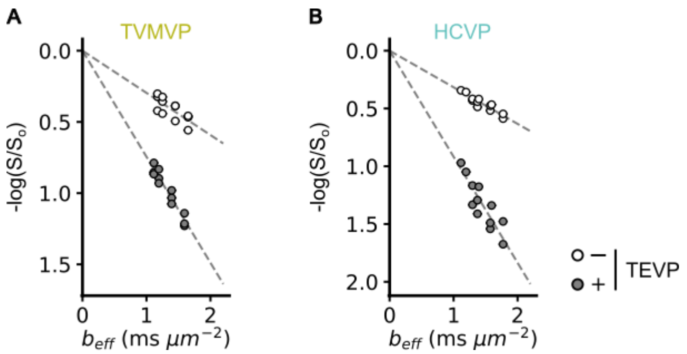


**Figure S12. Modularity of ER-URCAS**.

(A) Representative plots depicting the logarithmic decay in normalized diffusion-weighted signal intensity as a function of the diffusion-weighting factor ($b_{eff}$) in three different cell types expressing TVMVP-responsive ER-URCAS. The rate of diffusion (diffusivity) was determined from the slope of the decay curve

(B) Representative plots depicting the logarithmic decay in normalized diffusion-weighted signal intensity as a function of the diffusion-weighting factor ($b_{eff}$) in three different cell types expressing HCVP-responsive ER-URCAS. The rate of diffusion (diffusivity) was determined from the slope of the decay curve.

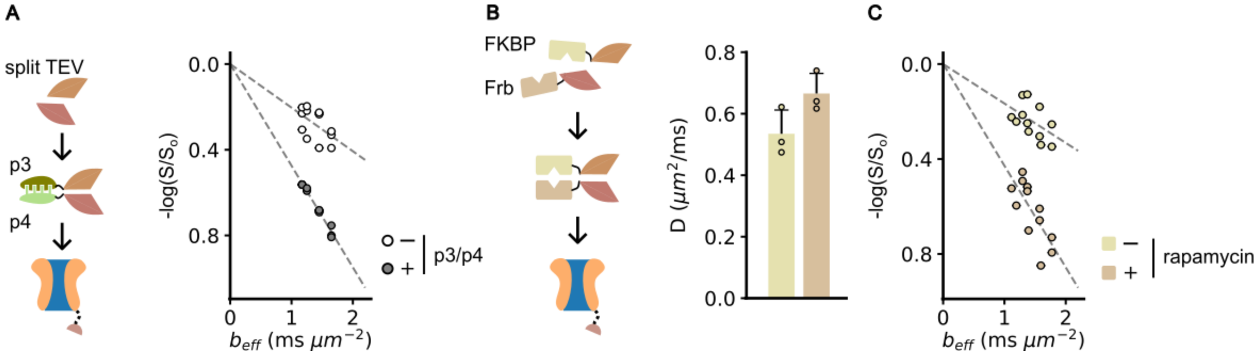


**Figure S13. Programmable biosensor design based on ER-URCAS**.

(A) Schematic illustrating the adaptation of ER-URCAS to detect biomolecular binding via reconstitution of split TEVP activity.

(B) Representative plots depicting the logarithmic decay in normalized diffusion-weighted signal intensity as a function of the diffusion-weighting factor ($b_{eff}$) in CHO cells engineered to express split TEVP-based ER-URCAS heterodimerization sensor. The rate of diffusion (diffusivity) was determined from the slope of the decay curve.

(C) Schematic illustrating the adaptation of ER-URCAS to detect chemically induced dimerization.

(D) Representative plots depicting the logarithmic decay in normalized diffusion-weighted signal intensity as a function of the diffusion-weighting factor ($b_{eff}$) in CHO cells engineered to express ER-URCAS sensor engineered to detect chemically induced dimerization. The rate of diffusion (diffusivity) was determined from the slope of the decay curve.

(E) Diffusivities of CHO cells expressing the split-TEVP-based ER-URCAS sensor to detect rapamycin-induced dimerization of FKBP and Frb. In these cells, TEVP contains activity-enhancing mutations, specifically S219V, and a truncation of C-terminal residues, which may contribute to elevated baseline diffusivity in the absence of rapamycin.

**Table 1. List of plasmids generated in this study**.

| Plasmid | Function |
|---|---|
| pANC12 | FLAG-hAqp1-TEVcs-ER50degron-IRES-EGFP |
| pANC13 | FLAG-hAqp1-TEVcs-FKBP12degron-IRES-EGFP |
| pANC14 | FKBP12degron-TEVcs-hAqp1- FLAG-IRES-EGFP |
| pANC21 | RPN4degron-TEVcs-hAqp1- FLAG-IRES-EGFP |
| pANC46 | FLAG7 hAqp1-TEVcs-FKBP12degron-IRES-EGFP |
| pANC47 | FLAG7 hAqp1-$G_4$S-TEVcs-FKBP12degron-IRES-EGFP |
| pANC48 | FLAG7 hAqp1-TEVcs-$G_4$S- FKBP12degron-IRES-EGFP |
| pANC49 | FLAG7 hAqp1-$G_4$S-TEVcs-$G_4$S- FKBP12degron-IRES-EGFP |
| pANC61 | FLAG7 hAqp1-GSG-(TEVcs-GSG)$_{3x}$-FKBP12degron-IRES-EGFP |
| pANC62 | FLAG7 hAqp1-($G_4$S)$_{3x}$-TEVcs-FKBP12degron-IRES-EGFP |
| pANC72 | FLAG7 hAqp1-GSG-(TEVcs-GSG)$_{2x}$-FKBP12degron-IRES-EGFP |
| pANC77 | FLAG7 hAqp1-GSG-(HCVcs-GSG)$_{3x}$-FKBP12degron-IRES-EGFP |
| pANC78 | FLAG7 hAqp1-GSG-(TVMVcs-GSG)$_{3x}$-FKBP12degron-IRES-EGFP |
| pANC82 | FKBP12degron-GSG-(TEVcs-GSG)$_{3x}$-hAqp1-FLAG-IRES-EGFP |
| pANC83 | RPN4degron-GSG-(TEVcs-GSG)$_{3x}$-hAqp1- FLAG-IRES-EGFP |
| pANC91 | RPN4degron-GSG-(TVMVcs-GSG)$_{3x}$-FLAG7hAqp1-GSG-(TEVcs-GSG)$_{3x}$-FKBP12degron-IRES-EGFP |
| pANC93 | FLAG7hAqp1-GSG-(TEVcs-GSG)$_{3x}$-FKBP12degron-IRES-uTEV3-t2A-EGFP |
| pANC56 | FLAG7 hAqp1-GSG-(TEVcs-GSG)$_{3x}$-KKYL-IRES-EGFP |
| pANC92 | FLAG7 hAqp1-GSG-(HCVcs-GSG)$_{3x}$-KKYL-IRES-EGFP |
| pANC99 | FLAG7 hAqp1-GSG-(HCVcs-GSG)$_{3x}$-KKYL-IRES-HCVP-t2A-EGFP |
| pANC100 | FLAG7 hAqp1-GSG-(TVMVcs-GSG)$_{3x}$-KKYL-IRES-EGFP |
| pANC101 | FLAG7 hAqp1-GSG-(TEVcs-GSG)$_{3x}$-KKYL-IRES-uTEV3-t2A-EGFP |
| pANC106 | FLAG7 hAqp1-GSG-TEVcs-GSG-KKYL-IRES-EGFP |
| pANC107 | FLAG7 hAqp1-GSG-(TEVcs-GSG)$_{2x}$-KKYL-IRES-EGFP |
| pNTA-uTEV3 | uTEV3-GSG-t2A-EBFP |
| pJW53 | uTEV3-DHFR2.0-IRES-mCherry |
| pANC75 | HCV NS3p-IRES-dTomato |
| pANC76 | TVMVp-IRES-dTomato |
| pANC79a | P4-GGGSGGG-cTEV-HA-IRES-dTomato |
| pANC79b | P3-Myc-nTEV-IRES-EBFP |
| pANC80a | cTEV-HA-IRES-dTomato |
| pANC80b | Myc-nTEV-IRES-EBFP |
| pANC98a | 3xFLAG-NES-FKBP12-(G4S)$_4$-nTEV-IRES-dTomato |
| pANC98b | 3xFLAG-NES-FRB-(G4S)$_4$-cTEV-IRES-EBFP |
| pANC98c | 3xFLAG-NES-FRB-(G4S)$_4$-cTEV(219)-IRES-EBFP |
| pKMD04 | CAM-cTEVp-t2A-nTEVp-RS20-IRES-mCherry |

**Table 2. Amino acid sequences of genetic parts used and developed in this study.**

| Plasmid | Sequence |
|---|---|
| FLAG | DYKDDDDK |
| Myc | EQKLISEEDL |
| HA | YPYDVPDYA |
| t2A | (GSG)EGRGSLLTCGDVEENPGP |
| ER tag | KKYL |
| hAqp1 | ASEFKKKLFWRAVVAEFLATTLFVFISIGSALGFKYPVGNNQTAVQDNVKVSLA FGLSIATLAQSVGHISGAHLNPAVTLGLLLSCQISIFRALMYIIAQCVGAIVAT AILSGITSSLTGNSLGRNDLADGVNSGQGLGIEIIGTLQLVLCVLATTDRRRRD LGGSAPLAIGLSVALGHLLAIDYTGCGINPARSFGSAVITHNFSNHWIFWVGPF IGGALAVLIYDFILAPRSSDLTDRVKVWTSGQVEEYDLDADDINSRVEMKPK |
| TEVcs | ENLYFQ\|S |
| TVMVp cs | ETVRFQ\|S |
| HCV NS3p cs | DEMEEC\|SQHL |
| ecDHFR degron | ISLIAALAVDRVIGMENAMPWNLPADLAWFKRNTLNKPVIMGRHTWESIGRPLP GRKNIILSSQPGTDDRVTRVKSVDEAIAACGDVPEIMVIGGGRVYEQFLPKAQK LYLSHIDAEVDGDTHFPDYEPDDWESVFSEFHDADALNSHSYCFEILERR |
| FKBP12 degron | GVQVETISPGDGRTFPKRGQTCVVHYTGMLEDGKKVDSSRDRNKPFKFMLGKQE VIRGWEEGVAQMSVGQRAKLTISPDYAYGATGHPGIIPPHATLVFDVELLKPE |
| Rpn4 degron | ASTELSLKRTLTDILEDELYHTNPGHSQFTSHYQNYHPNASITPYKLVNKNKEN NTFTWNHSLQHQNESSAASIPPQQT |
| ER50 degron | EFSLALSLTADQMVSALLDAEPPILYSEYDPTRPFSEASMMGLLTNLADRELVH MINWAKRVPGFVDLALHDQVHLLECAWMEILMIGLVWRSMEHPGKLLFAPNLLL DRNQGKCVEGGVEIFDMLLATSSRFRMMNLQGEEFVCLKSIILLNSGVYTFLSS TLKSLEEKDHIHRVLDKITDTLIHLMAKAGLTLQQQHQRLAQLLLILSHIRHMS SKRMEHLYSMKCKNVVPLSDLLLEMLDAHRL |
| TEVp (uTEV3) | GESLFKGPRDYNPISSTICHLTNESDGHTTSLYGIGFGPFIITNKHLFRRNNGT LLVQSLHGVFKVKNTTTLQQHLIDGRDMIIIRMPKDFPPFPQKLKFREPQREER ICLVTTNFQTKSMSSMVSDTSCTFPSSDGTFWKHWIQTKDGQCGNPLVSTRDGF IVGIHSASNFTNTNNYFASVPKNFMELLTNQEAQQWVSGWRLNADSVLWGGHKV FMVKPEEPFQPVKEATQLMN |
| TVMVp | GSSKALLKGVRDFNPISACVCLLENSSDGHSERLFGIGFGPYIIANQHLFRRNN GELTIKTMHGEFKVKNSTQLQMKPVEGRDIIVIKMAKDFPPFPQKLKFRQPTIK DRVCMVSTNFQQKSVSSLVSESSHIVHKEDTSFWQHWITTKDGQCGSPLVSIID GNILGIHSLTHTTNGSNYFVEFPEKFVATYLDAADGWCKNWKFNADKISWGSFT LVEDAPEDDFMAKKTVAAIMDS |
| HCV NS3p | GSGSVVLVGRLLLSGSGSAPITAYAQQTRGLLGCIITSLTGRDKNQAEGEVQIV STAAQTFLATCINGVCWTVYHGAGTRTIASPKGPVIQMYTNVDKDLVGWPAPQG TRSLTPCACGSSDLYLVTRHADVIPVRRRGDSRGSLLSPRPISYLKGSSGGPLL CPAGHAVGIFRAAVCTRGVAKAVDFIPVENLETTMRSPVFTDNSSPPAVS |
| P3 | SPEDEIQQLEEEIAQLEQKNAALKEKNQALKYG |
| P4 | SPEDKIAQLKQKIQALKQENQQLEEENAALEYG |
| nTEV | GESLFKGPRDYNPISSTICHLTNESDGHTTSLYGIGFGPFIITNKHLFRRNNGT LLVQSLHGVFKVKNTTTLQQHLIDGRDMIIIRMPKDFPPFPQKLKFREPQREER ICLVTTNFQT |
| cTEV | KSMSSMVSDTSCTFPSSDGIFWKHWIQTKDGQCGSPLVSTRDGFIVGIHSASNF TNTNNYFTSVPKNFMELLTNQEAQQWVSGWRLNADSVLWGGHKVFMSKPEEPFQ PVKEATQLMSELVYSQ |

| | |
|---|---|
| Calmodulin | DQLTEEQIAEFKEAFSLFDKDGDGTITTKELGTVMRSLGQNPTEAELQDMINEVDADGDGTIDFPEFLTMMARKMKDTDSEEEIREAFRVFDKDGNGYISAAELRHVMTNLGEKLTDEEVDEMIREADIDGDGQVNYEEFVQMMTAK |
| RS20 | RRKWNKTGHAVRAIGRLSS |
| nTEV | MGESLFKGPRDYNPISSTICHLTNESDGHTTSLYGIGFGPFIITNKHLFRRNNGTLLVQSLHGVFKVKNTTTLQQHLIDGRDMIIIRMPKDFPPFPQKLKFREPQREERICLVTTNFQ |
| cTEV | TKSMSSMVSDTSCTFPSSDGIFWKHWIQTKDGQCGSPLVSTRDGFIVGIHSASNFTNTNNYFTSVPKNFMELLTNQEAQQWVSGWRLNADSVLWGGHKVFMDKPEEPFQPVKEATQLMN |
| FKBP12 | GVQVETISPGDGRTFPKRGQTCVVHYTGMLEDGKKFDSSRDRNKPFKFMLGKQEVIRGWEEGVAQMSVGQRAKLTISPDYAYGATGHPGIIPPHATLVFDVELLKLE |
| FRB | LEMWHEGLEEASRLYFGERNVKGMFEVLEPLHAMMERGPQTLKETSFNQAYGRDLMEAQEWCRKYMKSGNVKDLLQAWDLYYHVFRRISK |
| nTEV | GESLFKGPRDYNPISSTICHLTNESDGHTTSLYGIGFGPFIITNKHLFRRNNGTLLVQSLHGVFKVKNTTTLQQHLIDGRDMIIIRMPKDFPPFPQKLKFREPQREERICLVTTNFQT |
| cTEV | KSMSSMVSDTSCTFPSSDGIFWKHWIQTKDGQCGSPLVSTRDGFIVGIHSASNFTNTNNYFTSVPKNFMELLTNQEAQQWVSGWRLNADSVLWGGHKVFM |

**References**


(1) Yun, J.; Huang, Y.; Miller, A. D. C.; Chang, B. L.; Baldini, L.; Dhanabalan, K. M.; Li, E.; Li, H.; Mukherjee, A. Destabilized Reporters for Background-Subtracted, Chemically-Gated, and Multiplexed Deep-Tissue Imaging. *Chem. Sci.* **2024**, *15* (28), 11108–11121. https://doi.org/10.1039/D4SC00377B.

(2) Nakahara, E.; Mullapudi, V.; Collier, G. E.; Joachimiak, L. A.; Hulleman, J. D. Development of a New DHFR-Based Destabilizing Domain with Enhanced Basal Turnover and Applicability in Mammalian Systems. *ACS Chem. Biol.* **2022**, *17* (10), 2877–2889. https://doi.org/10.1021/acschembio.2c00518.

(3) Chacko, A. N.; Miller, A. D. C.; Dhanabalan, K. M.; Mukherjee, A. Exploring the Potential of Water Channels for Developing Genetically Encoded Reporters and Biosensors for Diffusion-Weighted MRI. *J. Magn. Reson.* **2024**, *365*, 107743. https://doi.org/10.1016/j.jmr.2024.107743.

(4) Livak, K. J.; Schmittgen, T. D. Analysis of Relative Gene Expression Data Using Real-Time Quantitative PCR and the 2−ΔΔCT Method. *Methods* **2001**, *25* (4), 402–408. https://doi.org/10.1006/meth.2001.1262.

(5) Chowdhury, R.; Wan, J.; Gardier, R.; Rafael-Patino, J.; Thiran, J.-P.; Gibou, F.; Mukherjee, A. Molecular Imaging with Aquaporin-Based Reporter Genes: Quantitative Considerations from Monte Carlo Diffusion Simulations. *ACS Synth. Biol.* **2023**, *12* (10), 3041–3049. https://doi.org/10.1021/acssynbio.3c00372.

(6) Mukherjee, A.; Wu, D.; Davis, H. C.; Shapiro, M. G. Non-Invasive Imaging Using Reporter Genes Altering Cellular Water Permeability. *Nat. Commun.* **2016**, *7*, 13891. https://doi.org/10.1038/ncomms13891.
